# SubtiToolKit - a bioengineering kit for *Bacillus subtilis* and Gram-positive bacteria

**DOI:** 10.1101/2024.06.24.600474

**Authors:** Joaquin Caro-Astorga, Matt Rogan, Koray Malci, Hia Ming, Erika Debenedictis, Paul James, Tom Ellis

## Abstract

Building DNA constructs of increasing complexity is key to synthetic biology. Golden Gate methods led to the creation of cloning toolkits - collections of modular standardized DNA parts hosted on hierarchic plasmids, developed for yeast, plants, Gram-negative bacteria, and human cells. However, Gram-positive bacteria have been neglected. *Bacillus subtilis* is a Gram-positive model organism and a workhorse in the bioindustry. Here, we present the SubtiToolKit, a high-efficiency cloning toolkit for *B. subtilis* and Gram-positive bacteria. Its design permits DNA constructs for transcriptional units, operons, knock-in and knock-out applications. It contains libraries of promoters, RBSs, fluorescent proteins, protein tags, terminators, genome integration parts, a no-leakage genetic device to control the expression of toxic products during *E. coli* assembly, and a toolbox for industrially relevant strains of *Geobacillus* and *Parageobacillus* as an example of SubtiToolKit versatility for other Gram-positive bacteria and its future perspective as a reference toolkit.

## Introduction

Synthetic biology is demanding tools to speed up the assembly of complex genetic circuits. *Bacillus subtilis* is a Gram-positive bacteria that has been a mainstay in molecular biology and the bioindustry owing to its ease of genetic manipulability, strong protein expression and secretion capacity, and its regard as safe (GRAS) status (Souza et al., 2021; Yang et al., 2021).

Modern assembly methods for genetic engineering have leveraged the modularity of genetic parts to create complex genetic circuits(Rantasalo et al., 2018), combinatorial designs of biosynthetic pathways(Malcı et al., 2023), and perform precise genome edits(Malcı et al., 2022a). Golden Gate (GG) assembly is the most advanced cloning method, which is especially advantageous to create combinatorial libraries(Ellis et al., 2011; Engler and Marillonnet, 2014). The current bioengineering challenges call for a reliable assembly system and collections of standardized and characterized parts that can be exchanged between users. Since the development of the GG method in 2008, several toolkits have been developed with different designs in terms of structure, overhang syntax, tools, part libraries, or specificity for groups of organisms, with examples like MoClo for eukaryotic cells engineering (Weber et al., 2011), Yeast Toolkit(Lee et al., 2015; Malcı et al., 2022b), or EcoFlex designed for *E. coli*(Moore et al., 2016).

Briefly, GG utilises Type IIs restriction endonucleases cleave DNA outside their recognition sequences to generate 4bp overhangs which sequence can be designed to direct the order and orientation of DNA parts assembly. These restriction sites are smartly positioned in an inverse orientation flanking the genetic parts on their hosting vectors. Upon digestion of the hosting and destination vectors in a one-pot reaction, parts are released leaving the restriction site in the host vector and displaying overhangs directing the parts assembly. The restriction sites in the destination vector are inverted and released with the dropout marker gene. In the same pot, T4 ligase is added to seal the assembled fragments. Thermocycling the reaction to 37°C and 16°C permits digestion and ligation cycles. Reassemblies into the original vector keeps the restriction site and are released again in the next digestion cycle until parts rearrange into the desired construct, ready to be transferred into *E. coli*.

Increasing efforts at characterizing genetic regulatory elements such as promoters, ribosome-binding sequences, and terminators have yielded libraries of parts tailored to different organisms for regulating gene expression. This has also been targeted for *B. subtilis*(Guiziou et al., 2016; Liu et al., 2018; Xu et al., 2020; Yang et al., 2017). Nonetheless, there remain only a few examples of modular DNA assembly toolkits available for *B. subtilis* that provide partial or inefficient assembly options. SEVA siblings were designed to build constructions for double recombination, but the cargo must be previously assembled using classical methods(Radeck et al., 2017). ProUSER 2.0 is based on SEVA standards and it is more flexible in the assembly of plasmids including a promoter, but the cargo needs to be produced externally(Falkenberg et al., 2021). BacilloFlex is a toolkit based in EcoFlex that allows the assembly of transcriptional units(Wicke et al., 2017), but it uses inefficient overhangs, limited complexity of parts that can be assembled in number, positions and levels of hierarchy.

Despite the massive work done in *Bacillus* sp. for research and biotechnological applications, the field of Gram-positive bacteria has remained faithful to classical cloning methods. We have designed a highly efficient, modular, and hierarchical cloning system to assemble DNA fragments in plasmids in *E. coli* before transforming the final host. We expect SubtiToolKit to revolutionize the bioengineering of *B. subtilis*. This toolkit can be used for other Gram-positive bacteria by developing boxes with species specific part libraries. As an example, we have developed a GeoBox for *Geobacillus* and *Parageobacillus* sp. As it uses *E. coli* for assemblies, we expect the SubtiToolKit to be extensively used even for Gram-negative bacteria by the development of new boxes to harness the highly efficient design of this tool kit.

## Results

### Design of the SubtiToolKit (STK)

The STK is a Golden Gate (GG) assembly toolkit that is based on standardized modular and hierarchical assembly similar to other toolkits(Weber et al., 2011), but uses a different overhang syntax to achieve a much higher efficiency. Briefly, the STK assembly scheme allows for the directional assembly of 4-6 basic genetic parts from level 0 vectors (promoter, RBS, N-tag, CDS, C-tag, and terminator) into a level 1 destination vector. Up to four level 1 destination vectors can be assembled into Level 2, and up to four level 2 vectors can be assembled again into level 1 and so on with no limit of assemblies.

The STK contains 2 series of vectors: i) pSTK Level 0, 1, and 2 vectors which facilitate the cloning and assembly of the genetic parts and transcriptional units in *E. coli*; ii) pSTK-EXP shuttle vectors derived from pNW33N which are replicative in *Bacillus* sp., *Geobacillus* sp. and other Gram-positive bacteria (Figure 1A).

**Figure 1.**
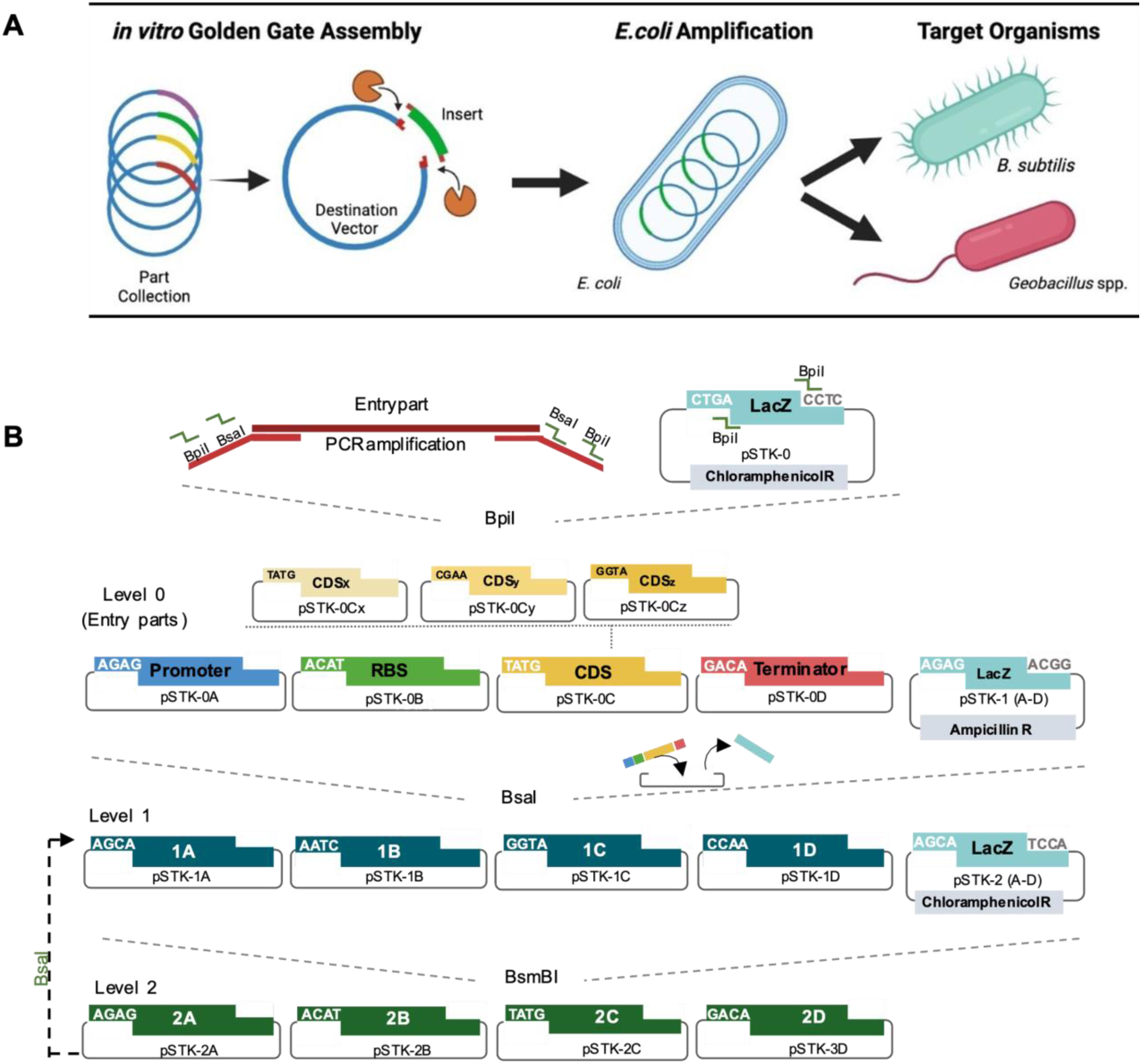
**The STK tool kit diagram**. A) STK workflow: parts are assembled *in vitro*, then transferred and amplified in *E. coli*, and then transferred to the final Gram-positive host organism. B) Entry parts are cloned from PCR using primers tailed with the sequence to be integrated into the entry plasmid pSTK0 by Golden Gate using BpiI. These primers also contain the BsaI site and the overhang syntax that determines the sequence position A-D once assembled into level 1 to produce a transcriptional unit (TU). Alternatively, C parts can be cloned into Cx-Cy- Cz to fuse N- and C- terminal tags to the CDS. TUs can be assembled in position 1A-1D, and up to 4 TUs can be assembled into any of the 2A-2D plasmids. Likewise, parts hosted in 2A-2D can be assembled into up to 16 TUs in any of the 1A-1D level plasmids and so on. As the cloning jumps through the levels, the system uses alternating restriction enzymes and antibiotics to select the correct assemblies.

The STK vectors pSTK Level 0, 1, and 2 vectors were derived from pBP of the EcoFlex^7^ toolkit with modifications. The STK vectors contain either a superfolder sfGFP or a lacZ-alpha fragment as a dropout marker to facilitate the screening of recombinant plasmids after GG assembly. The STK vectors adopt an optimized set of 4bp overhangs for high-fidelity GG reactions as reported by Potapov *et al*.(Potapov et al., 2018) to replace the fusion sites used in the EcoFlex toolkit, except for the TATG overhang used to ligate the RBSs to CDSs to preserve the ATG start codon within the overhang, reducing the assembly scar to one base pair to reduce any effect on expression levels. The pSTK assembly vectors contain a ColE1 high-copy number origin of replication for *E. coli* to maximize the yield along the assembly process. Plasmids of the same level contain the same antibiotic resistance and they alternate throughout the levels to facilitate the screening of correct assemblies into the correct destination vector (Figure 1B, and Supplementary Figure 1).

Level 0 vector pSTK-0 is the unique host for basic genetic parts such as promoters, RBSs, CDSs, protein tags, or terminators. Basic parts are incorporated by PCR amplification, or oligo annealing for short fragments (Figure 1B). The primers must contain a specific tail (Supplementary Table 1) with a BpiI site for Level 0 cloning and BsaI site with a specific overhang syntax that will determine the level 1 position as follows: A for promoters, B for RBSs, C for CDSs, and D for terminators. C position can be spitted in Cx for N-tags, Cy for CDSs, and Cz for C-tags (note that they use different overhangs and are not exchangeable with C parts). After PCR purification, a 10-cycle GG reaction using BpiI, pSTK-0, and the PCR product is used to transform *E. coli*. White colonies should be then confirmed by PCR. It is advisable to sequence the entry fragments for Level 0 to discard any mutation. For the next levels, only PCR is required to confirm the assembly as the probability of mutation is very low. The toolkit contains short spacers for every level and position to maximize assembly flexibility for the assembly of operons or other complex designs. An extra set of level 2 spacers contains a multi-cloning site AarI-SmaI/XmaI-BAmH-SphI-HindIII-Aarl to import/export fragments with classical cloning methods if required.

The assembly of Level 1 constructs from Level 0 basic parts can be hosted into four Level 1 destination vectors (pSTK-1A, -1B, -1C, -1D) using BsaI restriction enzyme. These plasmids have a different syntax to direct their assembly. Similarly, the assembly of Level 2 constructions from Level 1 sequences can be hosted into four Level 2 destination vectors (pSTK-2A, -2B, -2C, -2D) using BsmBI. Plasmids from level 2 syntax are designed to be assembled into any level 1 vectors again using BsaI (Figure 1B).

### Assembly validation and cloning efficiency in E. coli

To validate the assembly of the STK we assembled well-characterized parts in *E.coli* that allow a fast and easy evaluation with the naked eye. For Level 1, we constructed a transcriptional unit (TU) for GFPmut3b into a pSTK-1A vector (see methods). We tested the syntax assembly efficiency by a GG thermocycling protocol of 5-25 cycles. A thermocycling protocol of 10 cycles is shown to be sufficient for almost 100% assembly efficiency and dozens of colonies (Figure 2A). Increasing the number of cycles produced an increase in colony number. A 25-cycle protocol did not show any improvement compared to 20 cycles.

**Figure 2.**
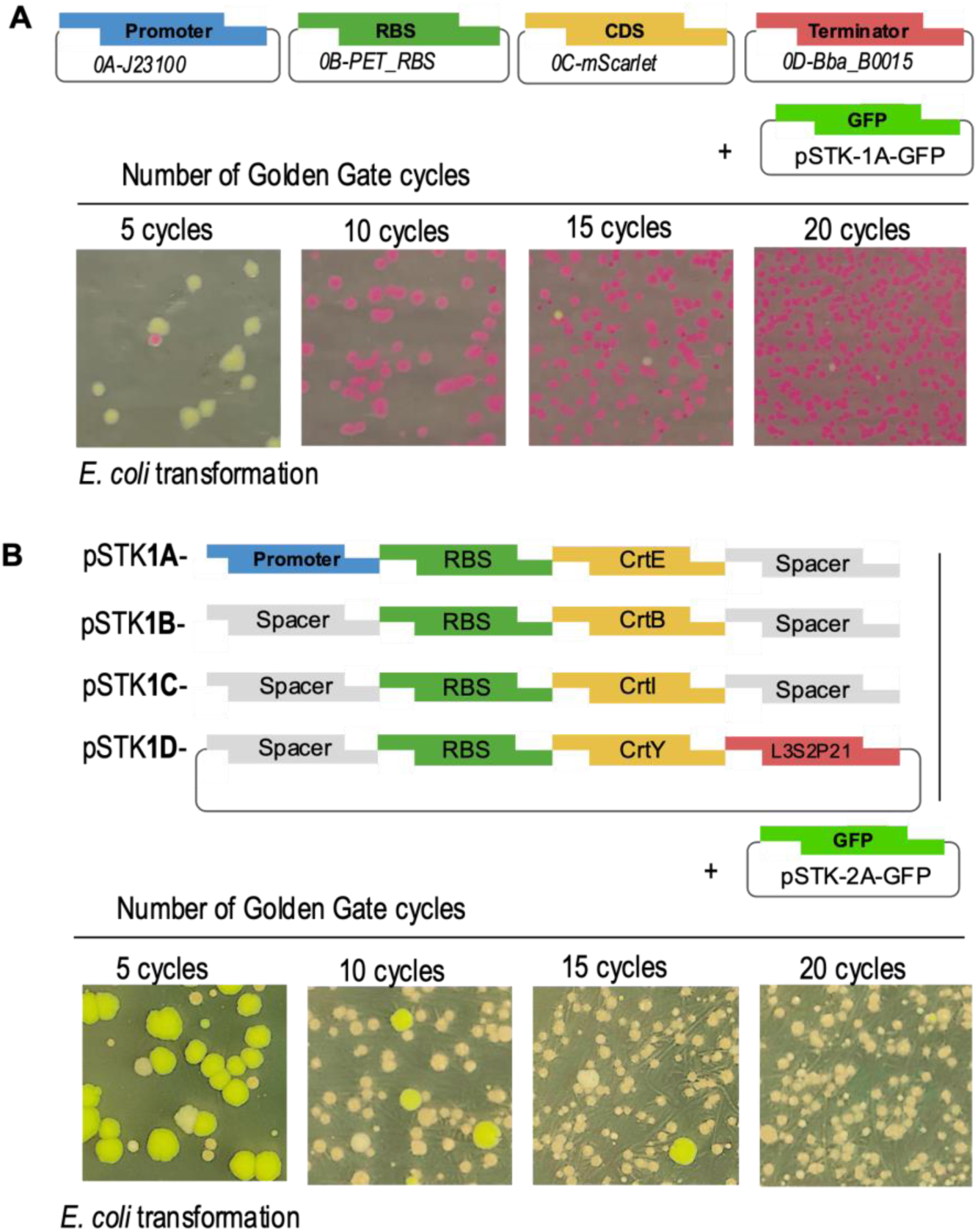
Assembly validation and efficiency of STK plasmids in *E. coli*. Level 1 assembly of a transcriptional unit to express mScarlet (A) and level 2 assembly of the carotene operon *ctrE/B/I/Y*. The high efficiency of the overhangs permits that 10 cycles of Golden Gate are enough to obtain dozens of colonies with a rate of correct assembly close to 100% after 10 or more cycles.

To validate level 2 assembly, we assembled an operon of 4 ORFs using the four genes *ctrE-B-I-Y* of the carotene operon(Nishizaki et al., 2007). To do so, the four ORFs were cloned into positions 0C-CDS and then into level 1 positions A-D. To build an operon, the promoter and the terminator were only assembled for the first and last gene respectively, using spacers for the internal 0A-promoter and 0D-terminator positions of the operon (Figure 2B). The four genes in level 1 A-D were then assembled to form the operon in a level 2 destination vector. The production of carotenes was confirmed by 1) colour, but sometimes it was difficult to differentiate, so we also used a method 2) colony-PCR of 10 colonies, and 3) the visual colour of a pellet from 5mlculture. We confirmed that all ten colonies were correct. The assembly efficiency of the level 2 syntax was close to 100 % (Figure 2C).

Generally, this high number of colonies yielded with that efficiency is not required for most applications. This means it is possible to reduce the use of enzyme and DNA plasmid and still obtain dozens of colonies, which means a reduction in the cost per assembly. We tested GG reactions serially diluted by 50% using T4 buffered 1x and also keeping the same DNA quantities but reducing the amount of enzymes in the same proportions. Results showed an acceptable reduction of efficiency to around 86.5% for using 50% reduction of enzymes independently of the DNA quantities. Higher dilutions showed a high drop in efficiency (Supplementary Figure 2). Reducing reaction volume to 5 µL is also feasible to keep concentrations, but there is a risk of evaporation during the thermocycling run.

### B. subtilis pEXP shuttle vector validation and efficiency

The basic STK plasmids used for parts assembly are not replicative in *B. subtilis*. The last assembly step before *B. subtilis* transformation should be performed in any of the two shuttle vectors pEXP, designed from the plasmid pNW33N (Figure 3). We domesticated it for the STK syntax and removed the incompatible restriction sites. It hosts a *repB* gene encoding for the rolling-circle replication initiator protein that allows plasmid replication in Gram-positive bacteria.

**Figure 3.**
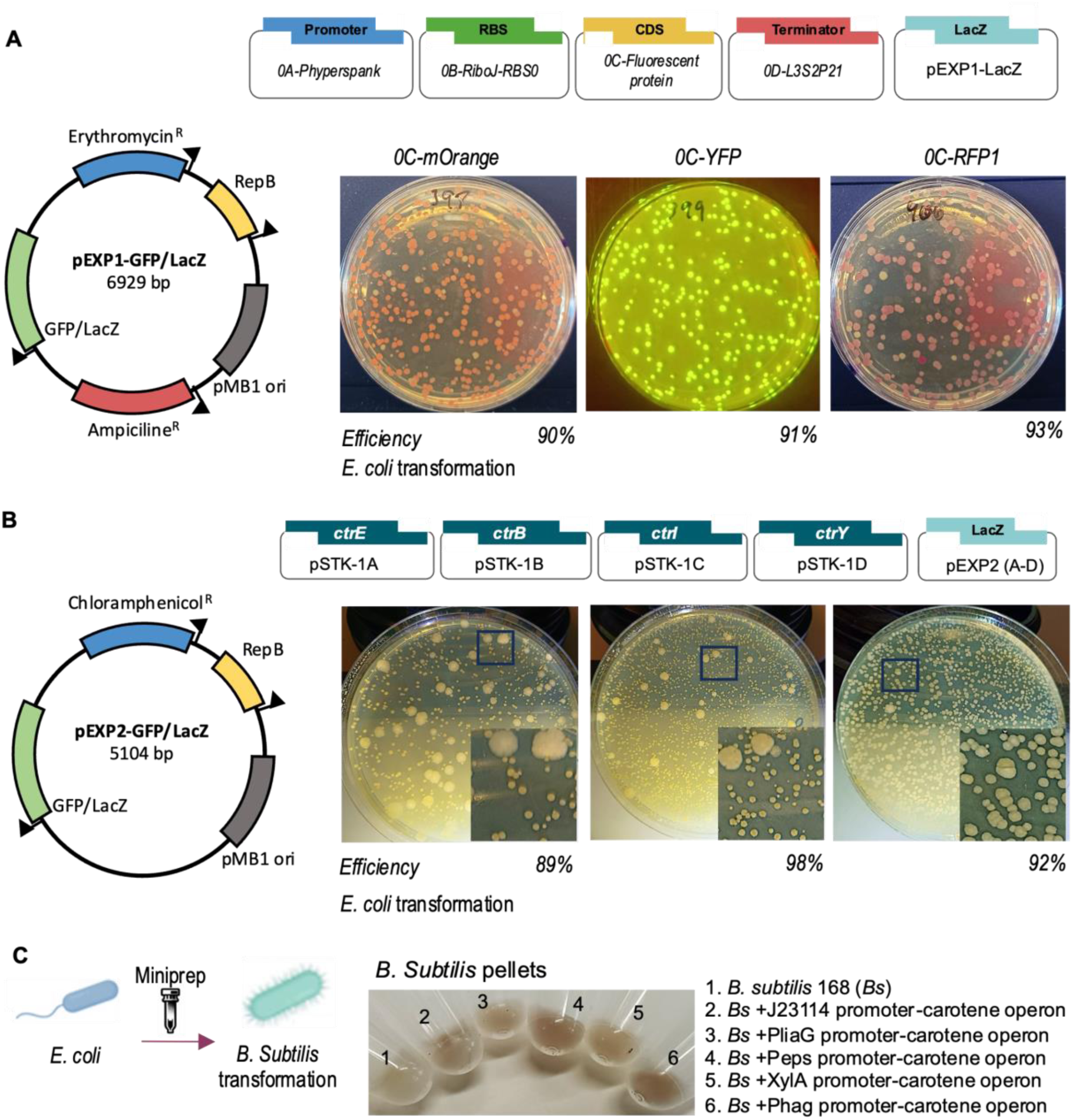
***B. subtilis* pEXP shuttle vectors assembly validation and efficiency**. (A) Genetic maps of level 1 pEXP-1 replicative plasmids for transcriptional unit expression and Golden Gate (GG) assembly of transcriptional units for mOrange, YFP and RFP1 fluorescent proteins to validate pEXP1 assembly. Images show agar plates after transformation of *E. coli* with GG reactions. (B) Genetic maps of pEXP2 for level 2 assemblies and GG assembly scheme of the carotene operon assembly. Image shows three replicates *E. coli* transformation of GG reactions. (C) Assemblies of the carotene operon under different promoters were assembled into pEXP2, transformed into *E. coli* and purified plasmids transformed into *B. subtilis* to confirm its stability. Image shows culture pellets of the different constructs. The integrity of the construction was also tested using PCR (Supplementary Figure 2F).

The assembly in *E. coli* was validated for pEXP level 1 with constructions for the expression of the fluorescent proteins GFPmut3b, eCFP, mCherry, mScarlet, mOrange, YFP, and RFP1 (Figure 3A) and in level 2 assembling the carotene operon (Figure 3B and Supplementary figure 3). These constructions were transferred to *B. subtilis* to confirm the plasmid stability. *B. subtilis* wild-type strain is not a good chassis for carotene expression as it produces a low amount of the precursor farnesyl diphosphate and it is mainly derived to another metabolic pathway producing farnesol(Filluelo et al., 2023). To test transformation, pellets of our strains expressing the carotene operon under different promoters showed differential intensities of a reddish colour compared to the white wild-type colonies. The integrity of the carotene operon in the plasmid was further confirmed by colony PCR (Figure 3C and Supplementary Figure 3F).

### Parts characterization in B. subtilis

Large libraries of native promoters(Liu et al., 2018) and RBSs(Guiziou et al., 2016) have been already characterized for *B. subtilis* and other Gram-positive species(Rondthaler et al., 2024). The STK contains a collection of 4 inducible and 9 constitutive promoters, 11 RBSs, and 12 terminators (Figure 4). These three parts determine the level of expression of the CDS they are flanking. To characterize promoters, we built assemblies using the promoter library and GFPmut3b as a reporter in *E. coli*, and then transformed into *B. subtilis* 168 to characterize their expression cultures in 24-well plates over 24 h in a plate reader where the fluorescence was monitored (Figure 4B). The control of the different inducible promoters is varied in terms of molecule of induction (mannitol, xylose, glycerol and mannose), growth phase (mid-log and stationery), leakage, maximum expression achieved and concentration of the inductor. We also characterized constitutive promoters, with 3P as the one that achieved highest expression (Figure 4C). We include Phyperspank and xylose promoters without lacI and xylR regulators to be used them for constitutive expression. The STK also contains a sporulation promoter sspB (Supplementary Figure 4). This provides a wide variety of options for bioengineers attending to desired outputs.

**Figure 4.**
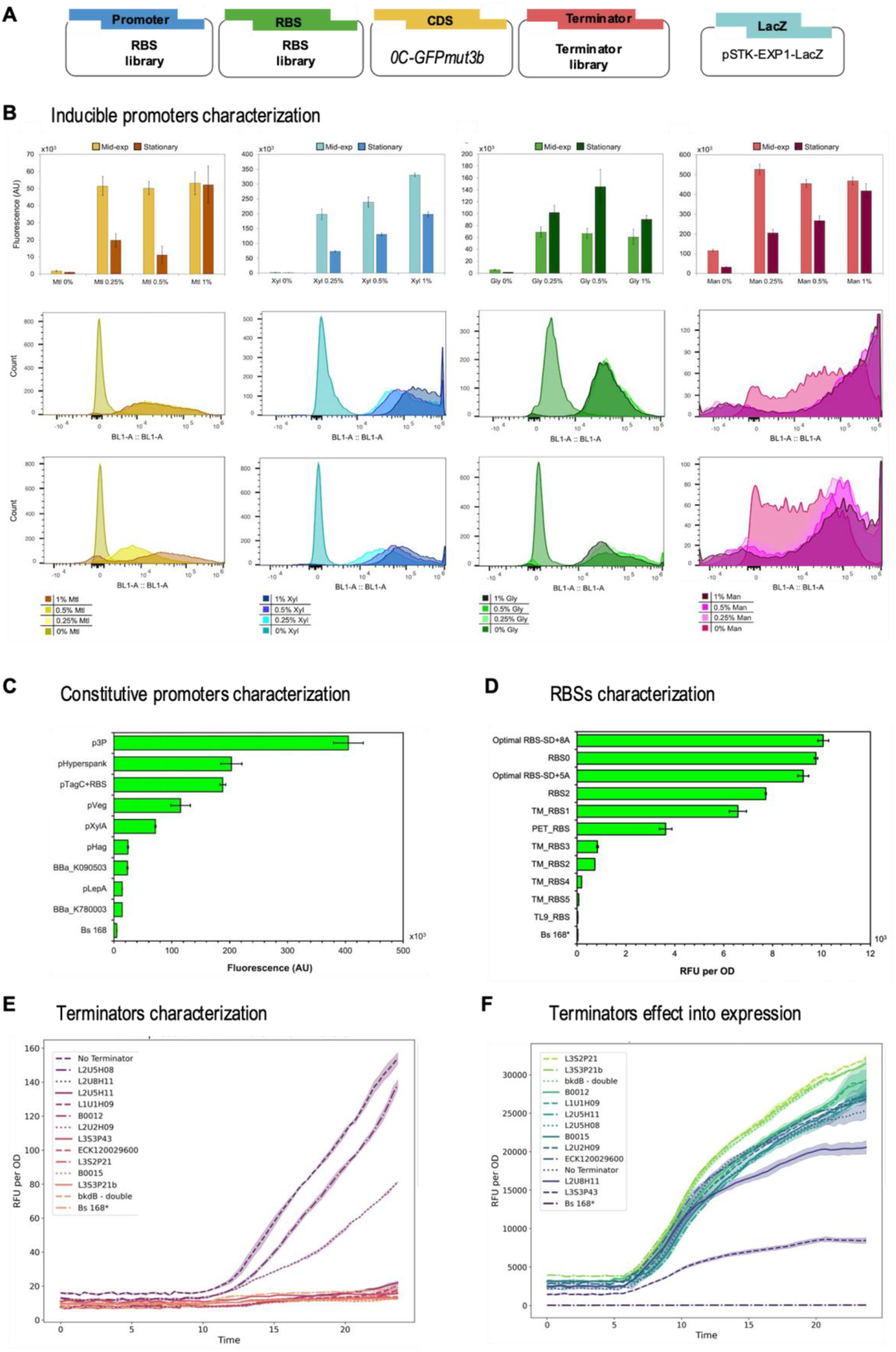
**Parts characterization in B. subtilis**. (A) Scheme of the Golden Gate (GG) assembly. (B) Characterization of the inducible promoters using GFPmut3b as a reporter gene: upper graphs of fluorescence intensity of GFPmut3b at mid- exponential stage (6 h) and stationary stage (24 h) after induction with different concentrations of mannitol (yellow), xylose (blue), glycerol (green), mannose (red). Below population distribution of expression measured in flow cytometry at mid- exponential phase at 6 h (middle graphs) and stationary phase at 24 h (lower graphs). (C) Characterization of constitutive promoters at stationary phase (24 h), using RBS TM_RBS3 and GFPmut3b as a reporter gene. The graphs show fluorescence measured in flow cytometry. (D) Characterization of RBSs using Phyperspank high expression promoter and GFPmut3b as a reporter gene. Tests with a low expression promoter were also characterized, showing low differences (Supplemerntary Figure 5). Bars represent the average fluorescence of three biological repeats, normalized by OD600, grown on 24-well plates in shaking conditions and 37°C, and measured in a plate reader. Error bars indicate SD. (E) RFP1 fluorescence graph for the characterization of terminators of a constructed operon with GFPmut3b and RFP1 interrupted with terminators placed between both CDSs. The graphs show red fluorescence (leakage) of the average of three biological repeats. The shade around the lines shows SD. (F) GFPmut3b fluorescence of constructions in E shows the effect of the terminators in protein expression. In E and F, legend is listed in order of values at the end of the experiment.

**Figure 5.**
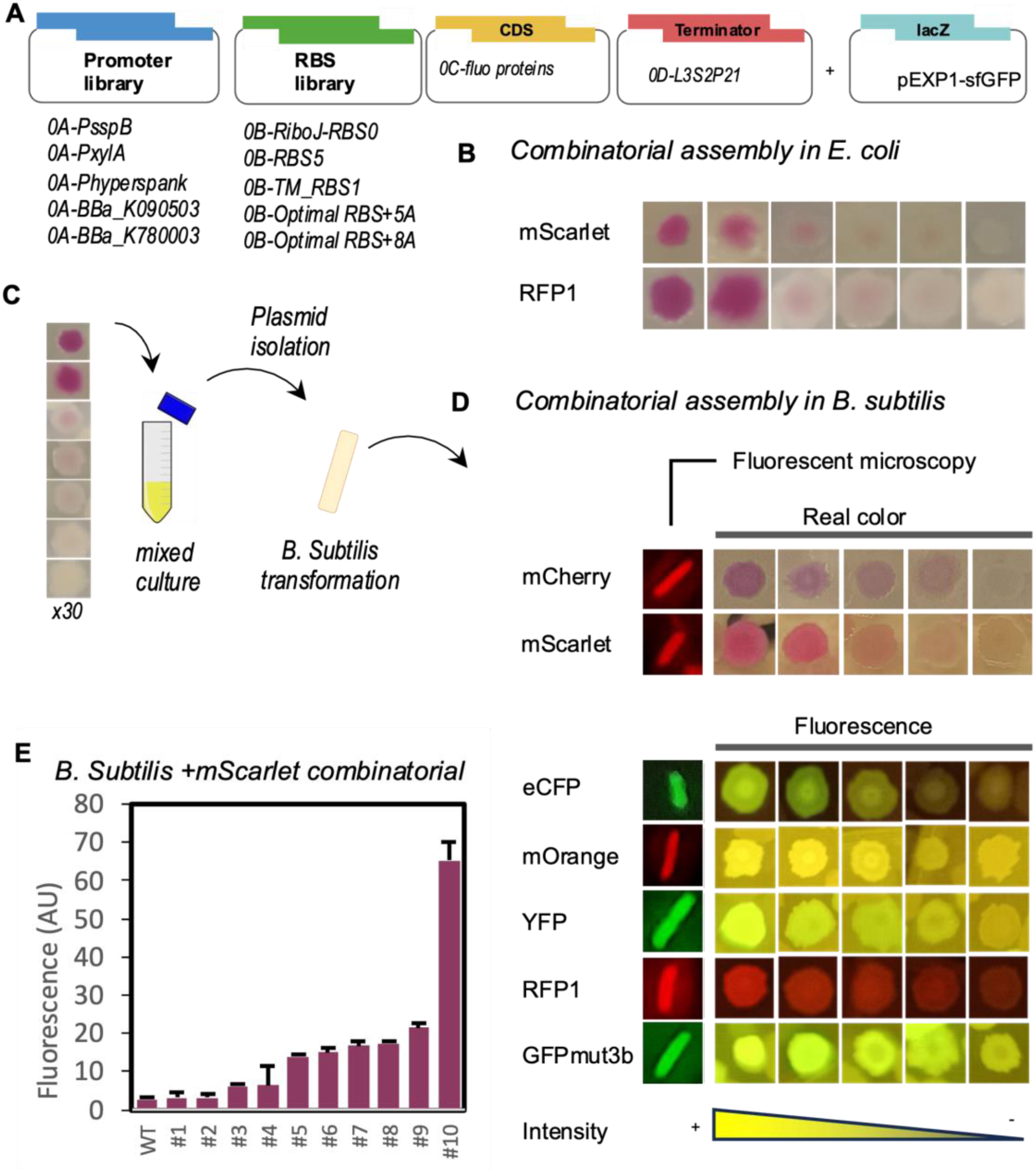
**Combinatorial assemblies**. (A) Scheme of the combinatorial assembly using libraries of promoters and RBSs. (B) Images of *E. coli* colonies showing a range of expression of mScarlet and RFP1 in real colour. C) Schematic of the process of combinatorial assemblies for *B. subtilis*. The colonies with the range of expression desired are selected and cultured together in a single culture tube from which miniprep can be done after a few hours of incubation and then transformed into *B. subtilis*. D) Images of *B. subtilis* colonies transformed with combinatorial assemblies of fluorescent proteins and fluorescent microscopy images of a single bacteria hosting the constructions. E) Characterization of RFP1 combinatorial assembly in *B. subtilis*.

Inducible promoters can be tuned by the concentration of the inducer. To modulate the expression of constitutive promoters, the STK contains a library of RBSs We observed that their relative behaviour among them can be different depending on the promoter used. We found that using a low expression promoter, all the RBSs provided a very similar fluorescent signal (Supplementary Figure 5) while using the high expression promoter Phyperspank showed a strong effect of the RBS in the outcome, covering the full range of expression levels (Figure 4D).

Terminators can also affect protein expression by affecting the stability of the mRNA or sensing chemicals that conditionally affect the efficiency of the terminator(Curran et al., 2015; F. De Felippes et al., 2020; Turnbough, 2019). We have characterized 12 terminators using a modified method by Gale *et al*. (Gale et al., 2021). Among the terminators tested, only L2U5H08 and L2U8H11 exhibited 75%-85% and 45%-55% leakage respectively, compared to the positive control (100% leakage) strain that had no terminator between GFPmut3b and mRFP1 (Figure 4E). The other terminators did not show a statistically significant difference in 24 h from the parent strain *B. subtilis* 168, which lacked the mRFP1 gene entirely. Notably, the bkdB-cold-double- terminator displayed no termination leakiness and significantly increased GFPmut3b expression due to mRNA stabilization, in agreement with previous characterizations(Welsch et al., 2015). Similar increases in expression were also observed with the terminators L3S2P21 and L3S3P21b (Figure 4F). This effect can be also used in combination with promoters and RBSs to further tune expression.

As a note for users, the behaviour of all these parts can be affected by the CDS present in the transcriptional unit, as it affects the secondary structure of the mRNA and its stability among other variables affecting expression. However, parts characterization in standard conditions is valuable as a guide for bioengineers.

### Combinatorial assemblies

As the final expression output of most genetic constructions are affected by the interaction of the parts, one of the most interesting advantages of genetic toolkits is the possibility of doing a single-pot-assembly of part libraries by adding a mix of promoters/RBSs/terminators together with a single ORF and a backbone as soon as it keeps the stoichiometry of DNA among positions. This produces all the possible combinations in the same pot, that can be transformed into *E. coli* and then colony screened for the desired output. This also reduces the necessity of a large library of promoters and RBSs in the toolkit given that the combination of a selection of a few of them would cover the entire range of expression. These combinatorial assemblies can produce hundreds or thousands of different constructions in a single GG reaction. The high efficiency of the STK is one of the most valued strengths for combinatorial assemblies as it drastically increases the coverage of combinations.

Transforming *E. coli* with a GG reaction can yield up to a few thousand colonies, while transforming *B. subtilis* directly is not efficient. Using a high-efficiency transformation method for *B. subtilis*(Vojcic et al., 2012), it is possible to obtain a few colonies with a 5x GG reaction (Supplementary Figure 3). This is a bottleneck for combinatorial assemblies. To overcome this limitation, we selected 35 different purple *E. coli* colonies from a GG reaction for the expression of every fluorescent protein included in the STK (Figure 4B) and mixed them in a 5mlculture, from which 3 μgr plasmid DNA was isolated and used to transform *B. subtilis* (Figure 4D). It showed the range of colonies fluorescence of each protein, and we also tested the fluorescence of all of them under fluorescent microscopy. For each assembly, many of the colonies obtained were slightly coloured. To get more insight into de degree of variety in expression we performed flow cytometry for mScarlet expression in *B. subtilis*, revealing differences in intensity by harbouring different promoter-RBS combinations (Figure 4E). Users should consider that eCFP and mOrange proteins can take 2 and 3 days respectively to maturate before been able to see coloured colonies.

### Genomic integration

It is particularly interesting for the industry to rely on genomic integration rather than plasmids that require antibiotic pressure to ensure they remain in the cells. Homologous recombination allows integrative vectors to be constructed by flanking genetic constructs with homologous regions (“homology arms”) upstream and downstream of the desired site of integration or to delete sections of the DNA. The thermosensitive shuttle vector pMAD was previously developed as a method to facilitate tractable genome integration of *B. subtilis*(Arnaud et al., 2004). This method requires plasmid curation after recombination by a temperature switch over several passages which sometimes turns into a tedious task. We designed STK versions of pMAD, but the efficiencies of assembly and plasmid stability were much lower than using STK plasmids - we speculate it is due to the big size of the plasmid. However, *B. subtilis* is much more efficient at taking linear DNA. In a GG reaction, we assembled 1000bp upstream and downstream homologous arms for recombination in *sacA* and *xynP loci.* We place these fragments in position 1A and 1D, flanking an arbitrary gene in position 1B and a spectinomycin cassette for recombinant selection in position 1C. The GG reaction excluded the destination vector to produce a linear DNA assembly. After transforming *B. subtilis* with a 5x GG reaction volume, it yielded hundreds of colonies (Supplementary Figure 7B). Approximately 50% of those colonies showed a correct integration when tested with colony PCR, and the other 50% probably contained the fragment expressing spectinomycin but only produced a single recombination or it was still floating in the cytoplasm (Supplementary Figure 7C). The STK contains homologous arms for the neutral *loci sacA, lacA and xynP*.

### Validation of N- and C- terminal assemblies

Protein engineering with C- and N- terminal tags is one of the basics in research and bioindustry labs for protein purification, secretion(Neef et al., 2021), increasing protein solubility, immunolocalization or protein-protein interaction, among many other applications. The STK allows to split the position 0-C for CDS into three positions named Cx, Cy and Cz to allocate tags at both sides - also using highly efficient overhangs. For N- terminal position, the STK contains a collection of seven *B. subtilis* native secretion tags (Apr, AmyQ, Cns, LipA, YlqB, SacB and TasA)(Guo et al., 2022). Secretion tags affect solubility, expression and secretion levels, requiring empirical trials with several secretion tags for each particular case. To validate the assembly, we fused these tags and a spacer as a control to amylase and galactosidase. However, *B. subtilis* secretion tags showed lethal toxicity in *E. coli* except for Csn that deeply affected growth, showing tiny blue colonies after 3 days of incubation, while using a spacer for 0-Cx position yielded a high number of blue colonies expressing galactosidase (Figure 6B and Supplementary Figure 8). This has been reported before for TasA secretion tag(Musik et al., 2021). To overcome this, we designed a 0B-RBS part containing a double terminator flanked by two recombination sites Bsdif specific for *B. subtilis* before the RBS(Sciochetti et al., 2001) (Figure 6C). This stops transcription in *E. coli*, permitting GG assembly (Figure 6D). Once transformed into *B. subtilis*, native recombinases RipX/CodV removes the double terminator and one recombination site, permitting transcription (Figure 6E). We built again constructions of galactosidase with secretion tags in *E. coli* and then transformed in *B. subtilis.* This device worked very well, but colony PCR showed that in some constructions the recombination did not happen in the entire pull of plasmid copies. We observed the loose of galactosidase activity over the passages, probably due to the burden of expressing *ganA* under a high expression promoter. This suggests that this device is more suitable for constructions integrated in the genome. Despite this handicap and also the high level of protease secreted by *B. subtilis* 168, secretion was high enough for Apr and SacB secretion tags to show green colour in the liquid culture when X-Gal was added (Supplementary Figure 8C). Nevertheless, we always recommend to test the non-classical protein secretion pathway of *B. subtilis* by using no tag, that can be much more efficient than the secretion tags as reported before and confirmed with the STK using galactosidase(Duan and Luan, 2023) (Supplementary figure 8C).

**Figure 6.**
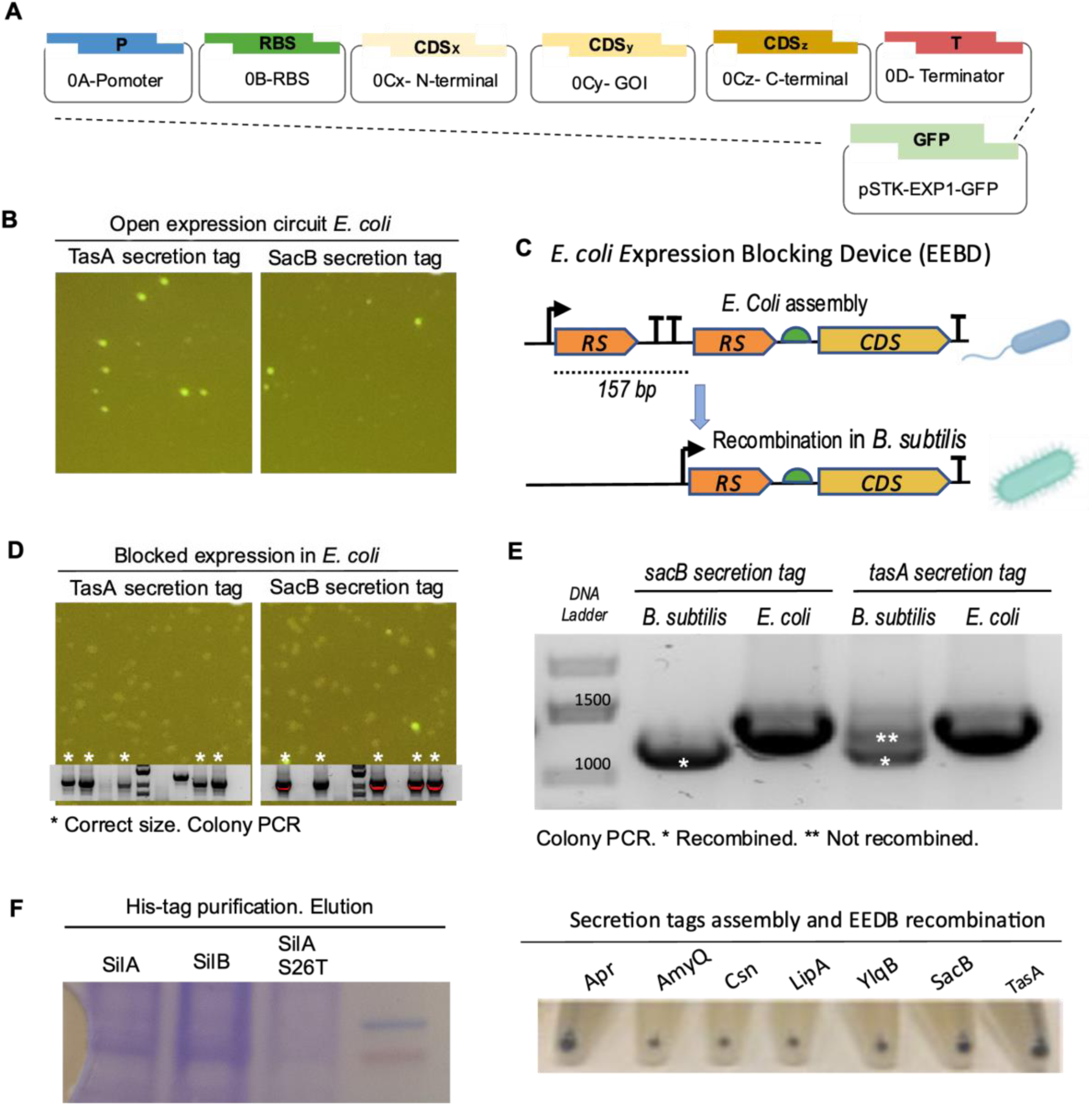
**Validation of N- and C- terminal assemblies**. (A) Scheme of the STK assembly of N- and C- terminal tags. (B) Amylase was fused to TasA and SacB secretion tags in a Golden Gate (GG) reaction. Images show the transformation in *E. coli* where the few white colonies were negative for PCR. (C) Genetic circuit scheme of an *E. coli* expression blocking device (EEBD) with a doble terminator between the promoter and the RBS, and flanked by two *B. subtilis-*specific recombination sites (RS). (D) Golden Gate of EEBD-amylase transformed into *E. coli* showed many white colonies positive for colony PCR (marked with an asterisk). (E) Agarose gel of colony PCR of *E. coli* and *B. subtilis* hosting the construction. In *B. subtilis*, the bands are lower indicating full recombination (SacB secretion tag) and partial recombination (TasA secretion tag) . Below, *B. subtilis* pellets transformed with EEBD-galactosidase with different secretion tags. (F) Acrylamide gel of three proteins fused to 6xHis tag at N- position and purified in cobalt columns.

The STK also contains C- terminal tags for protein purification (6xHis, FLAG, StrepII); SsrA to tune protein degradation; and SpyTag and SpyCatcher tags to induce covalent protein interactions. To validate this assembly position, we assemble 6xHis to three proteins and purified them in a cobalt column (Figure 6F). The STK also contains GFPmut3b in Cx and Cz positions for N- and C- terminal fusion. Spacers are also included when only one tag is desired.

### STK-GeoBox for Geobacillus sp

The Gram-positive bacterium *Geobacillus* and *Parageobacillus* are ideal candidates as microbial cell factories for sustainable production of industrially important chemicals due to their ability to grow at very high temperatures and can utilise a diverse range of substrates as an energy and carbon source, such as lignocellulose and crude oil(Hussein et al., 2015; Khaswal et al., 2022). Previous studies have expanded the available genetic tools for engineering different species of *Geobacillus*, which include vectors and libraries of basic parts(Bartosiak-Jentys et al., 2013; Koyama et al., 2022; Kurashiki et al., 2020; Madika et al., 2022; Pogrebnyakov et al., 2017; Reeve et al., 2016; Taylor et al., 2008). However, a toolkit for standardised modular and hierarchical assembly is not available. Despite the available tools, the characterization of these genetic parts has mostly been limited to *P. thermoglucosidasius* strains and omitting species with desirable phenotypes.

To address this, we have created an STK-GeoToolBox. The pS797-CCDB vector from Gilman *et al*. (2019) was used for the development of level 1 and level 2 GG destination vectors compatible with the STK(Gilman et al., 2019). The pS797-CCDB was based on the design of the pUCG18 vector, which contains a ColE1 and repBST1 origin of replication for plasmid replication in *E. coli* and *Geobacillus* respectively, a β-lactamase gene for ampicillin resistance, a TK101 thermostable kanamycin resistance gene and OriT for conjugal plasmid transformation, as well as a CCDB counter-selection marker with flanking inverse BsaI sites for level 1 assembly(Cripps et al., 2009; Gilman et al., 2019). To make a *Geobacillus* vector compatible with the STK, CCDB counter-selection marker was replaced with either a level 1 or level 2 LacZ or sf-GFP dropout, containing either inverse BsaI or BsmBI endonuclease sites respectively. Four pGeoEXP *Geobacillus* vectors were successfully made for the STK (pGeoEXP-sfGFP-1, pGeoEXP-sfGFP- 2, pGeoEXP-LacZ-1, pGeoEXP-LacZ-2).

We created an STK compatible library of promoters and characterised them under a strong RBS (PGEO79-RBS) in four species of *Geobacillus* (*P. thermoglucosidasius* DSM2542, *G. subterraneus* strain-K, *P. teobii* DSM14590T, *G. uzenensis* DSM13551)(Gilman et al., 2019). We included the ribozyme RiboJ upstream of the RBS to prevent incorporation of potential native 5’ untranslated regions from these promoters(Lou et al., 2012). Microplate reader analysis was used to characterise Dasher-GFP expression, with promoter strength normalised to Dasher-GFP concentration (nmol)/ OD600 (Figure 8A, Supplementary Figures 10 and 11). The resulting library displayed 96-fold, 165-fold, 238-fold, and 354-fold range libraries for *P. thermoglucosidasius*, *G. subterraneus*, *P. toebii,* and *G. uzenensis* respectively. As discussed before, parts may have different behaviours in different species even if they are close in evolution. Additionally, flow cytometry was performed to measure changes in expression across the cell population (Figure 8B).

**Figure 8.**
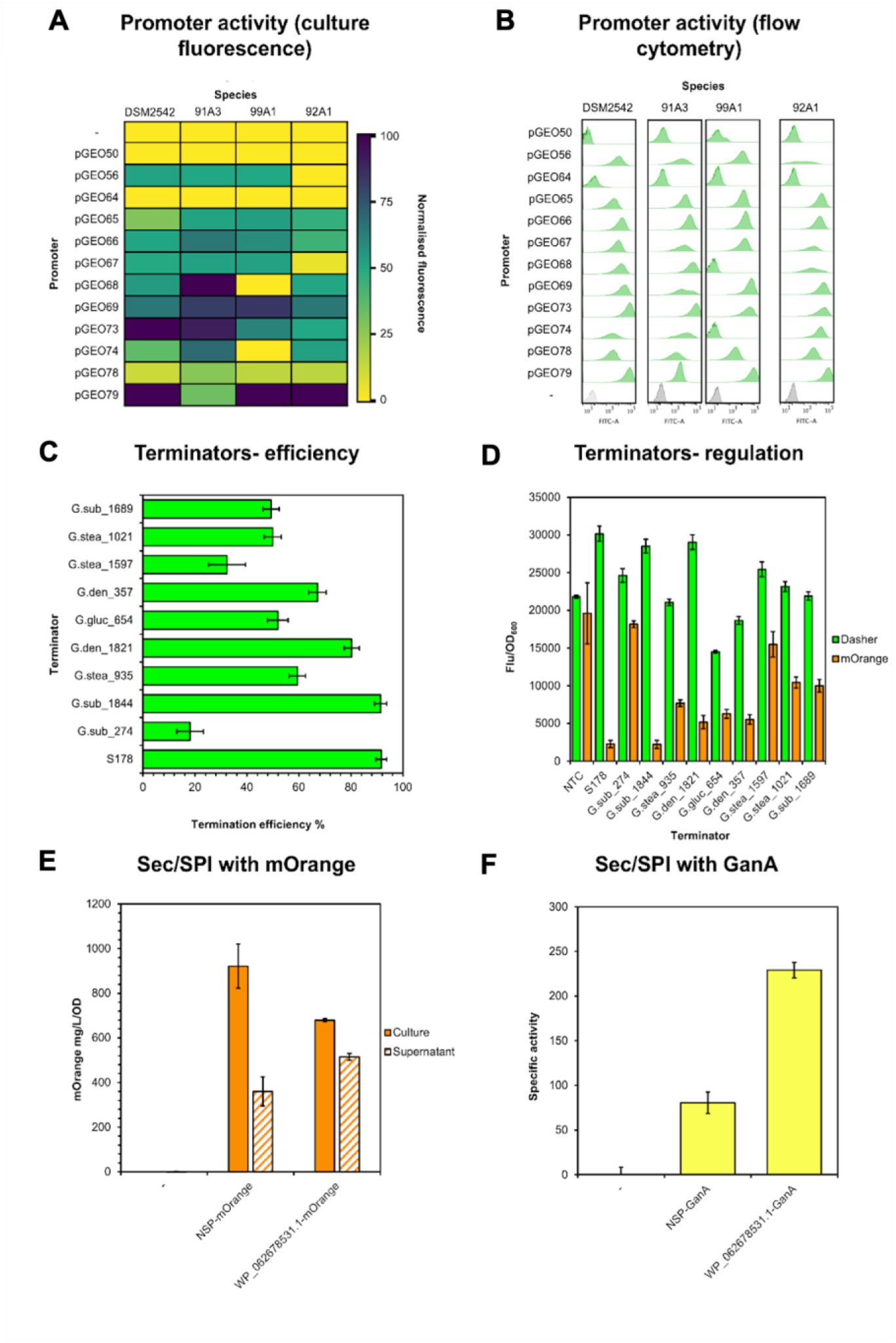
*STK-GeoBox for Geobacillus*. (A). Heatmap of constitutive promoters in four species of *Geobacillus* (DSM2542- *Parageobacillus thermoglucosidasius*, 91A3- *G. subterraneus* strain- K, 99A1- *P. teobii* DSM14590T, 92A1- *G. uzenensis* DSM13551). Promoters were fused to Dasher-GFP and characterised in each species after 16 h growth by measuring Dasher-GFP (nmol)/ OD600. Promoter strength was normalised to the strongest promoter in each species. An empty pGeoEXP control was used. (B). Flow cytometry characterization of constitutive *Geobacillus* promoters in different species after 16 h. The histogram shows the distribution in Dasher-GFP production. (C). Characterization of termination efficiency of native *Geobacillus* rho- independent terminators in *P. thermoglucosidasius*. Termination efficiency was calculated by analyzing the levels of Dasher-GFP upstream from the test terminator, to mOrange downstream from the test terminator and comparing this to a no-terminator control. Constructs were under the expression of PGEO79. (D). The impact of rho-independent terminators on upstream and downstream expression in *P. thermoglucosidasius* DSM2542. (E). Using the Sec/SPI signal peptide SPWP_062678531.1 for the secretion of mOrange *P. thermoglucosidasius* DSM2542. mOrange production in the cell culture and supernatant normalised to OD600. mOrange was under the expression of PGEO79. *P. thermoglucosidasius* DSM2542 with mOrange lacking a signal peptide and or an empty pGeoEXP vector was used as a negative control. (F). Using the Sec/SPI signal peptide SPWP_062678531.1 for the secretion of beta-galactosidase GanA from *Bacillus subtilis* 168 in *P. thermoglucosidasius* DSM2542 and characterization using ONPG assay. GanA without a signal peptide was used as a negative control. Error bars for all graphs represent the standard deviation of three biological repeats.

Ten rho-independent terminators native to the *Geobacillus* and *Parageobacillus* genus were characterised in *P. thermoglucosidasius* DSM2542 and three with either high, medium or low termination efficiency were included in the GeoToolBox. Terminators were identified using the ARNold database, with putative stem-loops, uracil tails, and free energy levels calculated. These terminators were characterised in *P. thermoglucosidasius* DSM2542 using a modified method by Gale *et al*. using Dasher-GFP and mOrange separated by the putative terminator(Gale et al., 2021; He et al., 2020). The termination efficiencies varied from 32.3% ± 5.1 to 93.1% ± 2.1 (Figure 8C). Additionally, effects of these rho-independent terminators on the expression of upstream CDS were also observed, with high Dasher-GFP production with the S178 terminator, and reduced Dasher-GFP expression levels with G.gluc_654, which could be their effect on mRNA stability (Figure 8D).

For targeted secretion of a heterologous protein, we included in the GeoToolBox the secretion tag Sec/SPI SPWP_062678531.1. Targeted protein secretion using SPWP_062678531.1 was validated in *P. thermoglucosidasius* DSM2542 by fusing the signal peptide with the fluorescent reporter mOrange and with GanA, which encodes a thermostable beta-galactosidase from *Bacillus subtilis* 168. Both CDSs were placed under constitutive expression. After 16 h growth, fluorescent measurements showed that 37.9% of mOrange was found in the supernatant for constructs lacking a signal peptide, compared to 77.7% found in the supernatant for mOrange with SPWP_062678531.1 (Figure 8E). Secretion was also determined for GanA by measuring the activity of the supernatant against ONPG. The results suggest that SPWP_062678531.1 results in increased beta- galactosidase activity in the supernatant compared to constructs lacking the signal peptide, with a 1.84-fold increase in activity compared to constructs lacking SPWP_062678531.1 (Figure 8F).

## Discussion

Since Golden Gate (GG) method was designed in 2008(Engler and Marillonnet, 2014), many toolkits have been developed for different groups of organisms and different designs were proposed, with this method achieving a quite mature stage(Bird et al., 2022). Nevertheless, there are still some aspects that can be modified for a better performance. We developed the SubtiToolKit (STK) for four reasons: 1) to cover a gap in GG tools for Gram-positive bacteria; 2) to design a hierarchy of plasmids that allows every cloning purpose, 3) to improve the efficiency of previous GG toolkits, and 4) to reduce the time and cost of the technique by using efficient overhangs syntax that require a shorter protocol and less amount of enzymes and DNA, based in the studies of *Potapov et al*.(Potapov et al., 2018; Pryor et al., 2020). As a result, the efficiency of the assembly of STK is around 99% after 10 GG cycles when accurately adjusted molarity of DNA parts is used, with low contamination of genomic DNA and correct adjustment of DNA concentrations. To facilitate these calculations, we provide a useful spreadsheet (Supplementary Spreadsheet file). This high efficiency allows much shorter protocols, saving up to 66% of the time if we consider a 30 cycles protocol as the generally accepted practice compared to 10 cycles required for STK. When required, a further number of cycles and volume of the reaction increases the number of correct constructs yielded in the pot. We also demonstrate that it is possible to transform a GG reaction directly into *B. subtilis*. This permits to avoid the *E.coli* transformation, selection, culture, and miniprep steps before transforming *B. subtilis*.

The STK has been designed to fit all cloning purposes - multiple TU expression, operons, N- and C-terminal tags, deletions and genomic integration. The fact that level 2 assemblies can be assembled into Level 1 positions permits unlimited size or constructions. The current limitation would be the stability of plasmids. As the field of synthetic biology demands bigger constructs, a cosmid can be domesticated into STK syntax.

We expect STK to be a reference for Gram-positive bacteria and also the research community to contribute to the further development of the STK by creating new tools, modules, gates, genetic devices, and boxes for other industrial strains of Gram-positive bacteria like *Streptomyces* sp., *Lactococcus* sp.*, Lactobacillus* sp. *or Clostridium* sp. Nonetheless, STK is already designed and validated for *E. coli* as a working strain. Just by domesticating other collections of parts into STK syntax an STK-ColiBox or boxes for other Gram-negative will quickly expand STK use to harness its advantages. As an example of its versatility for organisms other than *B. subtilis*, we have the STK-GeoBox for engineering *Geobacillus* and *Parageobacillus* species. Existing plasmids for specific bacteria can easily be modified to be compatible with the STK through the addition of compatible dropouts and ensuring no other endonuclease sites exist. In the same way, the STK syntax is not compatible with other tool kits, but parts can be easily domesticated by PCR. Despite new boxes being expected to contain specific parts from the target organisms, Gram-positive bacteria show a high transferability of parts between the groups of bacteria(Rondthaler et al., 2024). However, these parts need to be characterized in the new host as it is not rare that parts behave differently in different species, even from the same genus as we show in our results on different species of *Geobacillus* and *Parageobacillus* (Figure 6A).

Despite the benefits that STK offers, there is still space for improvement. The main limitation of the STK is the assembly of level 2 constructs which use BsmBI restriction enzyme. BsmBI has only 25% of activity in T4 ligase buffer required for GG and requires longer times of digestions, using 6-8 minutes of digestion to achieve complete digestion and the high efficiency that the STK syntax provides. We expect a better BsmBI enzyme to be developed in the near future to reduce this time and make level 2 assemblies as fast as level 1.

## STAR Methods

### Strains and Culture Conditions

*Escherichia coli* NEB Turbo and *Bacillus subtilis* 168 (BS168) strains were routinely cultured in LB agar or liquid, shaking at 250rpm, at 37°C, supplemented with appropriate antibiotics: ampicillin (100 µg/mL), chloramphenicol (20 µg/mL) or kanamycin (50 µg/mL) for *E. coli.* For *B. subtilis*, we used chloramphenicol (20 µg/mL), kanamycin (7.5 µg/mL) or erythromycin/lincomycin (1 µg/mL/25 µg/mL). For Blue-White screening, plates were supplemented with 0.02% X-Gal.

*Parageobacillus thermoglucosidasius* DSM2542, *Parageobacillus toebii* DSM14590T (99A1), *Geobacillus uzenensis* DSM13551 (92A1) and *Geobacillus subterraneus* strain-K (91A3) were grown in modified Lennox LB containing 0.59 M MgSO4, 0.91 M CaCl2, 0.04 M FeSO4 and 1.05 M nitrilotriacetic acid trisodium salt. For transformation, *Geobacillus* and *Parageobacillus* strains were selected using 12.5 µg mL^-1^ of kanamycin. All strains of *Geobacillus* and *Parageobacillus* were obtained from the *Bacillus* Genetic Stock Centre (Columbus, Ohio, USA). For conjugal plasmid transformation, *E. coli* S17-1 (genotype: *recA pro hsdR RP4-2-Tc::Mu-Km::Tn7*) was used as a donor host.

### DNA Manipulation and Plasmid Preparation

General PCR and restriction enzyme digestion and ligation were performed using reagents from New England Biolabs (NEB). PCR amplification for cloning was carried out using Q5® High-Fidelity DNA Polymerase, while Phire Plant Direct PCR Mastermix (ThermoFisher Scientific) was used for colony PCR screening. DNA purification from agarose gels was performed using ZymocleanTM Gel DNA Recovery Kit (Zymogen). Plasmid isolation was performed using QIAprep® Miniprep (Qiagen). Genomic DNA Purification was carried out using QIAGEN Genomic-tip 100/G (Qiagen). All procedures were carried out according to the manufacturer’s instructions.

### E. coli Transformation

Chemically competent *E. coli* NEB Turbo cells and *E. coli* S17-1 were used. 50µL of competent cells were incubated with DNA at 4°C for 10 minutes, followed by heat shock at 42°C for 1 minute, then cooled to 4°C for 1 minute and incubated at 37°C for 1 h. The cells were subsequently seeded onto LB agar plates with appropriate antibiotic selection.

### Bacillus Transformation

Chemically competent *B. subtilis* 168 cells were produced using the method developed by *Vojcic et al.*(Vojcic et al., 2012). This protocol produced highly competent cells that can be stored at -80°C keeping its competence.

### Geobacillus and Parageobacillus Transformation

A modified version of the protocol by Tominaga et al. (2016)(Tominaga et al., 2016) was used for the transformation of all *Geobacillus* and *Parageobacillus* species. Transformed *E. coli* S17-1 was used as donor cells for the transformation of *Geobacillus* and *Parageobacillus* recipient cells. A single *E. coli* S17-1 colony was collected and grown in 5mlLB without antibiotics and a single *Geobacillus* or *Parageobacillus* colony was grown in 10mlof mLB until an OD600 of 0.3 was reached for each. 1mlof *E. coli* culture was mixed with 9mlof *Geobacillus* or *Parageobacillus* culture, centrifuged at 4500 rpm for 5 minutes, the supernatant removed, and the pellet resuspended in 30 µl of LB. This resuspension was then placed onto an LB plate and incubated at 37°C for 16 h. After this, *E. coli* donor cells were incubated at 55°C for 1 h to kill *E. coli* donor cells. The mating mixture was resuspended in 200 µl of LB, and this mixture was plated out on a mLB/Kan (12.5 µg ml^-1^) and left 24-48 h at 55°C.

### Construction of SubtiToolKit Vectors

The STK vectors were generated from EcoFlex toolkit(Moore et al., 2016). The plasmids pBP and pTU1-A-RFP were linearised by PCR amplification using primers flanking the entry sites to remove the BsaI and BsmBI restriction enzyme recognition sites, overhangs, and RFP dropout cassettes. The drop-out sfGFP and LacZ cassettes were amplified from pKTK137(Goosens et al., 2021) and pBP-lacZ-alpha fragment respectively using primers with overhangs containing the appropriate BpiI, BsaI, and BsmBI restriction enzyme recognition sites and STK overhang syntax, as well as homology regions to the corresponding linearized plasmids to allow the final assembly by Gibson method (Supplementary Figure 1). A significant modification to the destination plasmids from the EcoFlex toolkit is the adoption of an optimized set of 4bp overhangs for high-fidelity Golden Gate assembly (∼99% fidelity) to replace the overhangs used in the EcoFlex standard, except for the TATG overhang used for part 0C to preserve the start codon ATG. Additionally, the BpiI restriction site was incorporated into the level 0 plasmids flanking the dropout cassettes to standardize and simplify the construction of basic part plasmids, instead of using the NdeI/NcoI and BamHI/BglII restriction sites as in the EcoFlex standard.

### Golden Gate Assembly

For Level 0, a 10 µl Golden Gate (GG) reaction containing 50 fmol of the pSTK-0-sfGFP vector and 50 fmol of the adapted basic part were mixed with 0.5 µl (10 units) of BpiI and 0.5 µl (200 units) of T4 ligase in 1X T4 ligase buffer. The reaction mixture was cycled 10-25 times at 37°C for 2 min and 16°C for 5 min. The reaction mixture was then incubated at 60°C for 2 min and 80°C for 5 min to denature the enzymes and was directly used to transform *E. coli* and plated on LB agar plates with 20 µg/mL chloramphenicol. Successful transformants were indicated by non- fluorescent colonies.

For GG assembly of Level 1 constructs, 0.5 µl (10 units) of BsaI were used and bacteria were plated in LB supplemented with ampicillin 100 µg/ml. For GG assembly of Level 2 constructs, 0.5 µl (10 units) of BsmBI were used and the digestion time was increased up to 8 minutes to overcome the reduced activity (25%) of BsmBI in T4 ligase buffer. Transformed *E.coli* were plated in LB supplemented with chloramphenicol 20 µg/ml.

### Parts characterization for STK

To characterize the dynamic behaviour of the STK parts, saturated overnight cultures of each construct were diluted 100 times in 1mlLB cultures with the appropriate antibiotics in clear, flat- bottom 24-well plates. The plates were incubated in a Tecan Spark® microplate reader for 24 hours, with double orbital shaking at 270 RPM and a temperature of 37°C. Measurements were taken every 30 minutes. Optical density (OD) was measured at 600 nm in absorbance mode. For fluorescence measurements, the excitation wavelengths were set to 485 nm and 560 nm, and the emission wavelengths to 520 nm and 620 nm for GFPmut3b and mRFP1, respectively, in bottom- reading mode. The manual gain was set to 50, and the z-position was calculated based on the first well (A1). The bottom surfaces of the lids facing the wells were covered with an anticondensation solution containing 0.05% Triton X-100 (Sigma-Aldrich) in 20% ethanol to prevent condensation on the lids.

For flow cytometry analysis, cultures were diluted 1/100 into phosphate-buffered saline (PBS) to a final volume of 200 µL in a 96-well plate. Measurements were taken using an Attune NxT Acoustic Focusing Cytometer with an autosampler module. Voltage settings were 440 V for FSC, 340 V for SSC, and 490 V for BL1 to measure GFPmut3b expression. A total of 10,000 events were recorded per sample. Singlets were gated using FSC-H x FSC-A. Data from the flow cytometer were analysed and visualized using FlowJo 10.10.0. Samples were collected at the 6th hour of cultivation, representing the mid-exponential phase, and at the 24th hour, representing the stationary phase. For characterizing inducible promoters, inducers were added when the cultures reached the early stationary phase (approximately two hours after inoculation). The experiments were conducted in triplicates.

### Parts characterization for STK-GeoBox

For characterization of promoters with Dasher-GFP in different species of *Geobacillus* and *Parageobacillus*, strains were plated on mLB/Kan, and individual colonies were picked and grown in 5mlmLB/Kan for 16 h at 55°C 200 rpm. Three individual colonies were selected for each strain, giving three independent biological repeats. 200 µl of culture was taken for microplate reader analysis using a clear, flat bottom 96-well plate. A Tecan Spark microplate reader was used to measure the optical density at 600 nm, and Dasher-GFP fluorescence was measured at 477 nm excitation and 522 nm emission (gain = 44, Z-position = 19131 µm). A 1/100 dilution of the culture was performed into phosphate buffer saline to a total of 300 µl. 100 µl was used for flow cytometry analysis using a BD FACSCanto™ II. The Voltage settings were as follows: 100 V FSC, 300 V SSC, 300 V FITC-A (all logarithmic). The threshold was set to 1,000 for FSC. A total of 100,000 events were measured per sample, giving 300,000 measurements per construct. Flow cytometry Data analysis was performed using FlowJo 10.9.0.

Terminators were characterised using constructs that contained Dasher-GFP and mOrange were characterised using microplate reader analysis. Test constructs were grown as described in 2.2.8.1 and fluorescence of Dasher-GFP was measured at 477 nm excitation and 522 nm emission (gain = 44, Z-position = 30,702 µm) and mOrange was measured at 588 nm excitation and 610 nm emission (gain = 111, Z-position = 30,702 µm). A no terminator control was used to determine the maximum amount of Dasher-GFP to mOrange without a terminator when used with the PGEO79 promoter. The termination efficiency was calculated using the method described by Gale et al. (38).

Signal peptide characterization was performed in *P. thermoglucosidasius* DSM2542, with reporters expressed using PGEO79 promoter and 50K_RBS (RiboJ upstream), with the Sec/SPI SPWP_062678531.1. For mOrange characterisation, a 40mlmLB/Kan culture was inoculated with 400 µl overnight and grown for 16 h at 55°C 200 rpm. Three biological repeats were made per construct. For culture fluorescence analysis, 200 µl was used. For mOrange supernatant analysis, 1mlof culture was centrifuged at 5000 x *g* for 10 min. The supernatant was filtered and 200 µl was used for plate reader analysis using Tecan Spark microplate reader to measure OD at 600 nm, and mOrange fluorescence was measured at (gain = 101, Z-position = 31,001 µm).

To determine the secretion of an active beta-galactosidase, an o-nitrophenyl-ß-d- galactopyranoside (ONPG) assay was performed. *P. thermoglucosidasius* with a pGeoEXP expressing GanA from *Bacillus subtilis* 168 with and without a Sec signal peptide was grown for 16 h in 40mlmLB/Kan at 55°C, 200 rpm. Cultures were then centrifuged at 5000 *x g* for 10 min and filtered to remove any cell debris using a 0.2 µm nitrocellulose membrane. The culture supernatant was concentrated 100 x using a 10 KDa Merck Millipore protein concentrator and the protein concentration was determined using a Qubit™ Protein Broad Range (BR) Assay Kits. Beta-galactosidase assays were performed in triplicate in microplates, containing 10 mg l^-1^ of total protein supernatant, 50 µl 0.06 M Na2HPO4-7H2O, 0.02 M NaH2PO4-H2O, 0.01 M KCl, 0.001 M MgSO4-7H2O (pH 7.0), and 17 µl of 4 mg/ml ONPG. After 30 min incubation at 50°C, 600 rpm, the reaction was stopped by adding 125 µl of 1 M sodium carbonate. This gives a final reaction volume of 192 µl. Absorbance at 420 nm was measured to determine how much ONPG was hydrolysed. The amount of ONPG hydrolysis was calculated using the following equation:

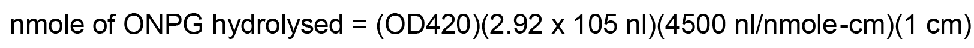

where OD420 is the absorbance, 1.92 x 10^5^ nl is the volume of the reaction and 4500 is the extinction coefficient. The specific activity of the culture supernatant was then calculated using the following equation:

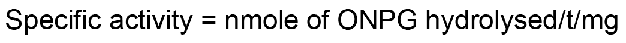

Where *t* is the time in minutes and *mg* is the amount of protein used in the reaction.

## Acknowledgments

The authors wish to thank Dr Chris French, Dra Alicia Clement, Dr Andre Pulsen, for their technical support and advice; Giovanni Manresa di Serracapriola, Shirrin Bamezai, Alessandro Serafini and Imeperial College Sporadicate iGEM Tam for their comments on the experience on using the toolkit ; Fangkang Meng, for his feedback using the STK for automation; and Dra Sheilla Ingemann Jensen for sharing ProUser2.0 plasmids.

This work has been founded by: JCA was funded by the US Office of Naval Research Global (ONRG) and US Army CCDC DEVCOM grant W911NF-18-1-0387 and UK Engineering and Physical Sciences Research Council (EPSRC) grant EP/N026489/1. JCA and ED were supported by the Francis Crick Institute which receives its core funding from Cancer Research UK (CC2239), the UK Medical Research Council (CC2239), and the Wellcome Trust (CC2239), and a Steel Perlot Early Investigator Grant. TE and KM were supported by the NextSkins EIC project, funded by the European Union’s Horizon Europe research and innovation programme under grant agreement number 101071159. MR was supported by a studentship obtained through the Hub for Biotechnology in the Built Environment which was an E3 Project from Research England.

## Authors contribution

JCA proposed the research, designed the toolkit, performed most of *B. subitilis* experiments and supervised HM and KM. HM performed cloning of most basic parts and STK plasmids and characterized level 1 assemblies. JCA and KM performed *B. subtilis* parts characterization. KM also contributed with cloning and assemblies. PJ supervised *Geobacillus* work, and TE supervised most of the experimental work in *B. subtilis*. JCA, KM, MR, PJ, and TE discussed the results and commented on the paper. ED commented on the paper and contributed to management. JCA coordinated the team.

This work was enabled due to contributions of the facilities of the Imperial College Centre for Synthetic Biology, The Francis Crick Institute and Northumbria University.

## Declaration of Interest

The authors declare no competing interests.

---

**Supplementary figure 1.**
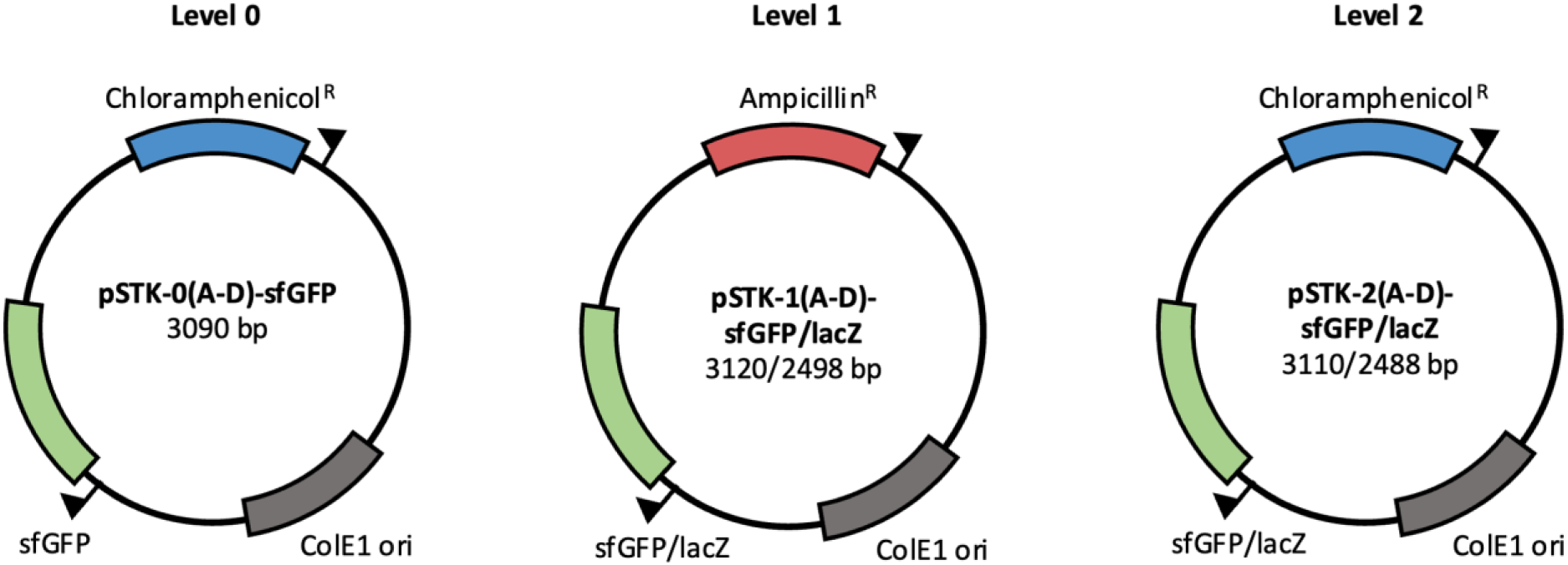
**SubtiToolKit (STK) plasmid maps**. Maps of the hierarchical plasmids used for parts assembly in *E. coli*. There is only one entry vector version with a sfGFP dropout gene as entry parts are not expressed. STK contains two versions of plasmids for levels 1 and 2 assemblies using sfGFP or lacZ-alpha fragment as the desired construction may contain fluorescent proteins and requires a different marker.

**Supplementary figure 2.**
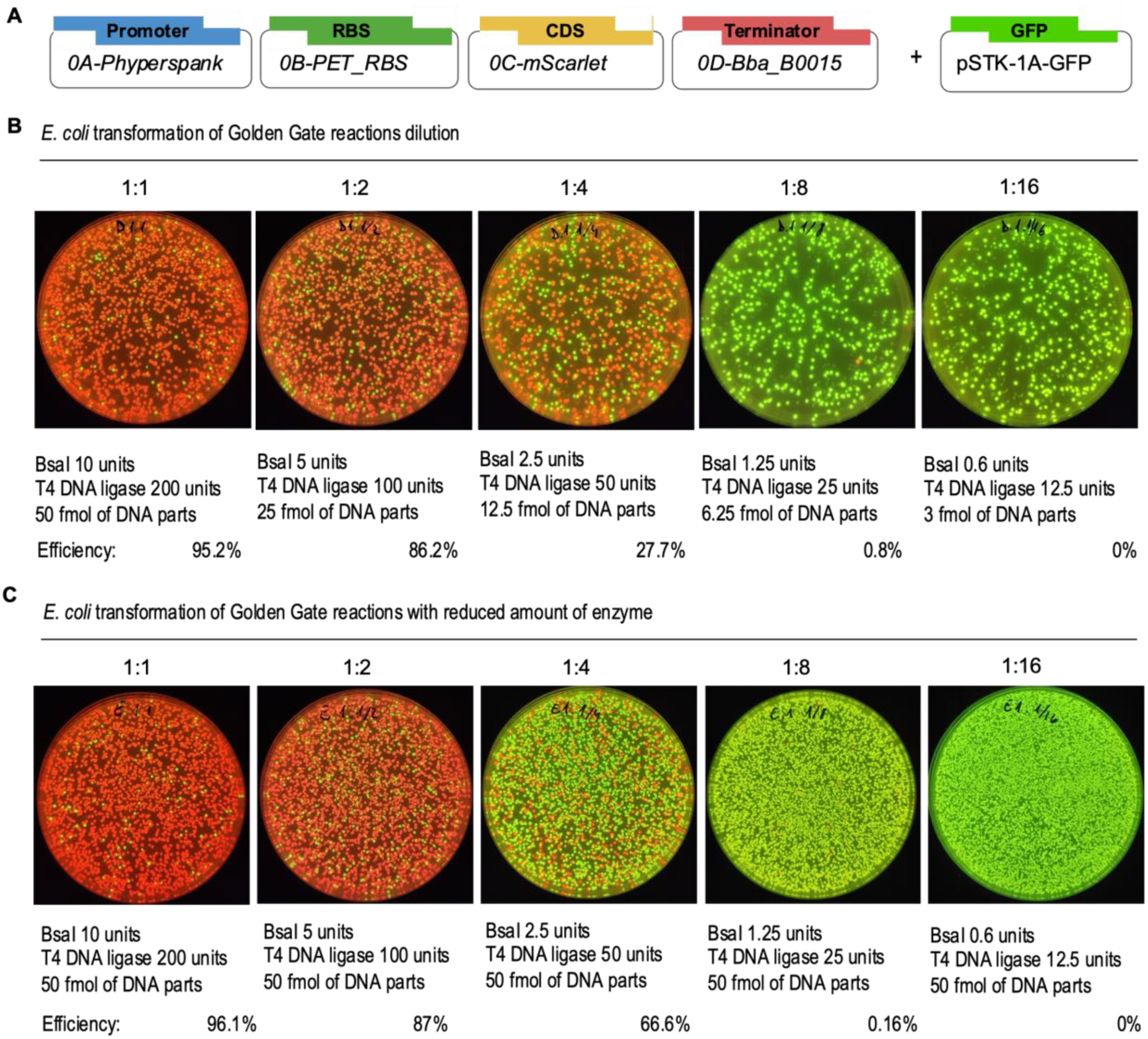
**Golden Gate efficiencies with reduced quantity of enzymes**. A) Scheme of the Golden Gate assembly parts used. B) A mix of Golden Gate reaction containing 10 units of BsaI, 200 units of T4 DNA ligase and 50 fmol of each part was serially diluted by 50% with T4 ligase buffer 1x. The reaction was run in a 20 cycles reaction protocol. Images show *E. coli* transformation of 10 μL of each reaction. Image representative of two replicates. C) Experiment repeated reducing only the quantity of enzymes and maintaining 50 fmols of DNA of each part.

**Supplementary Figure 3.**
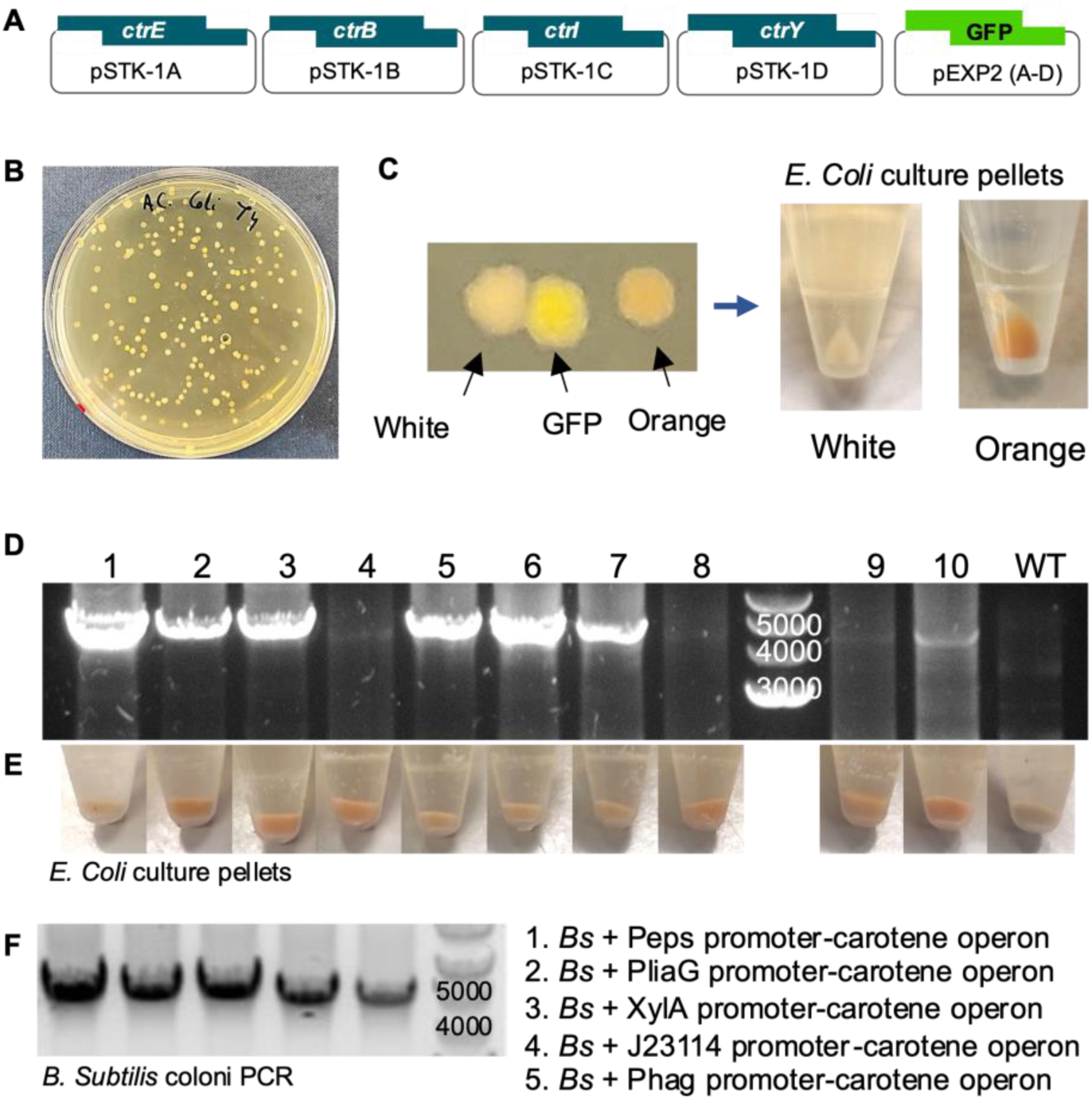
**Level 2 assembly validation**. (A) Schematic of the Golden Gate (GG) assembly. (B) Agar plate of an *E. coli* transformation of the GG reaction. (C) Images of the three types of colonies obtained. White (partial assembly) and fluorescent colonies (destination vector reassembled) are wrong assemblies which are hard to differentiate from orange colonies expressing the operon due to the yellow background of the agar plate. When colonies are picked into liquid cultures, pellets of cells show a characteristic orange colour. (D) Electrophoresis of colony PCRs of ten orange-ish *E. coli* colonies and (E) pellets of colony cultures. (F) Colony PCR of *B. subtilis* hosting the carotene operon under different promoters.

**Supplementary Figure 4.**
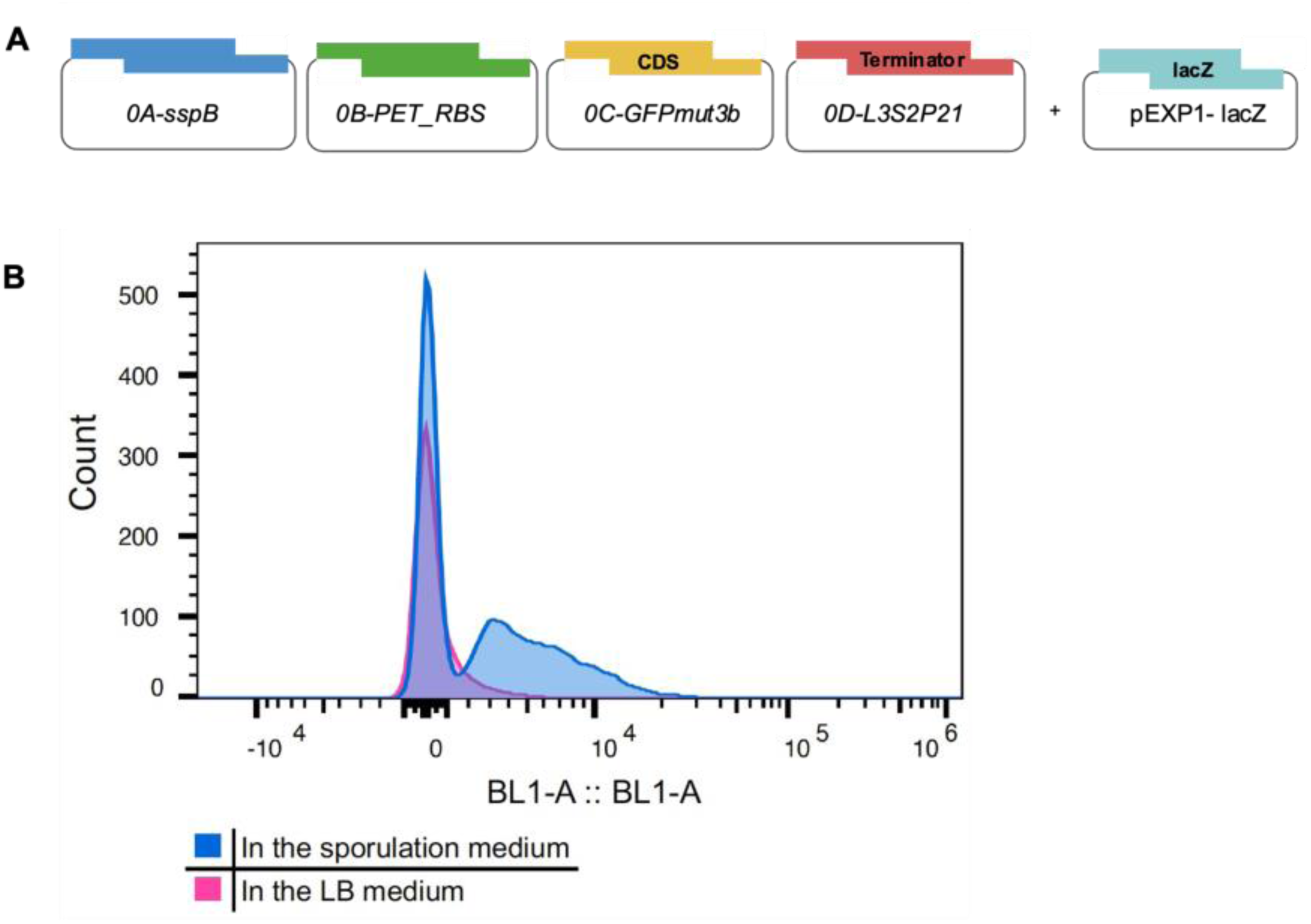
**Sporulation promoter characterization**. (A) Scheme of the Golden Gate assembly. (B) Characterization of the sporulation promoter sspB controlling GFPmut3B expression by fluorescence activity of bacteria cultures in sporulation media (blue) compared to LB media (magenta).

**Supplementary Figure 5.**
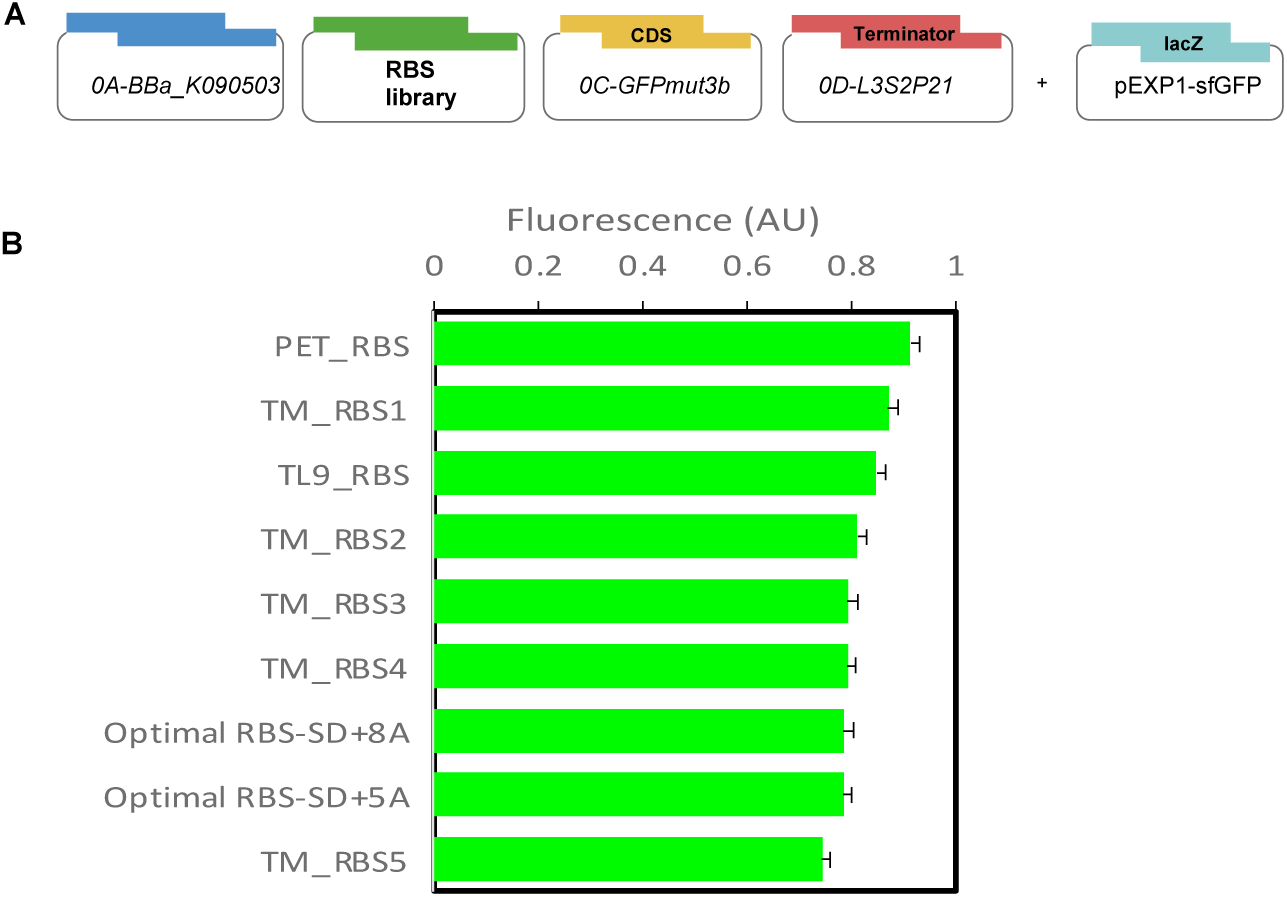
**RBSs characterization**. (A) Scheme of the Golden Gate assembly of the RBSs library. (B) The combination of a weak promoter with RBSs show poor differences between RBSs. Error bars indicate standard deviation of the average of three replicates.

**Supplementary Figure 6.**
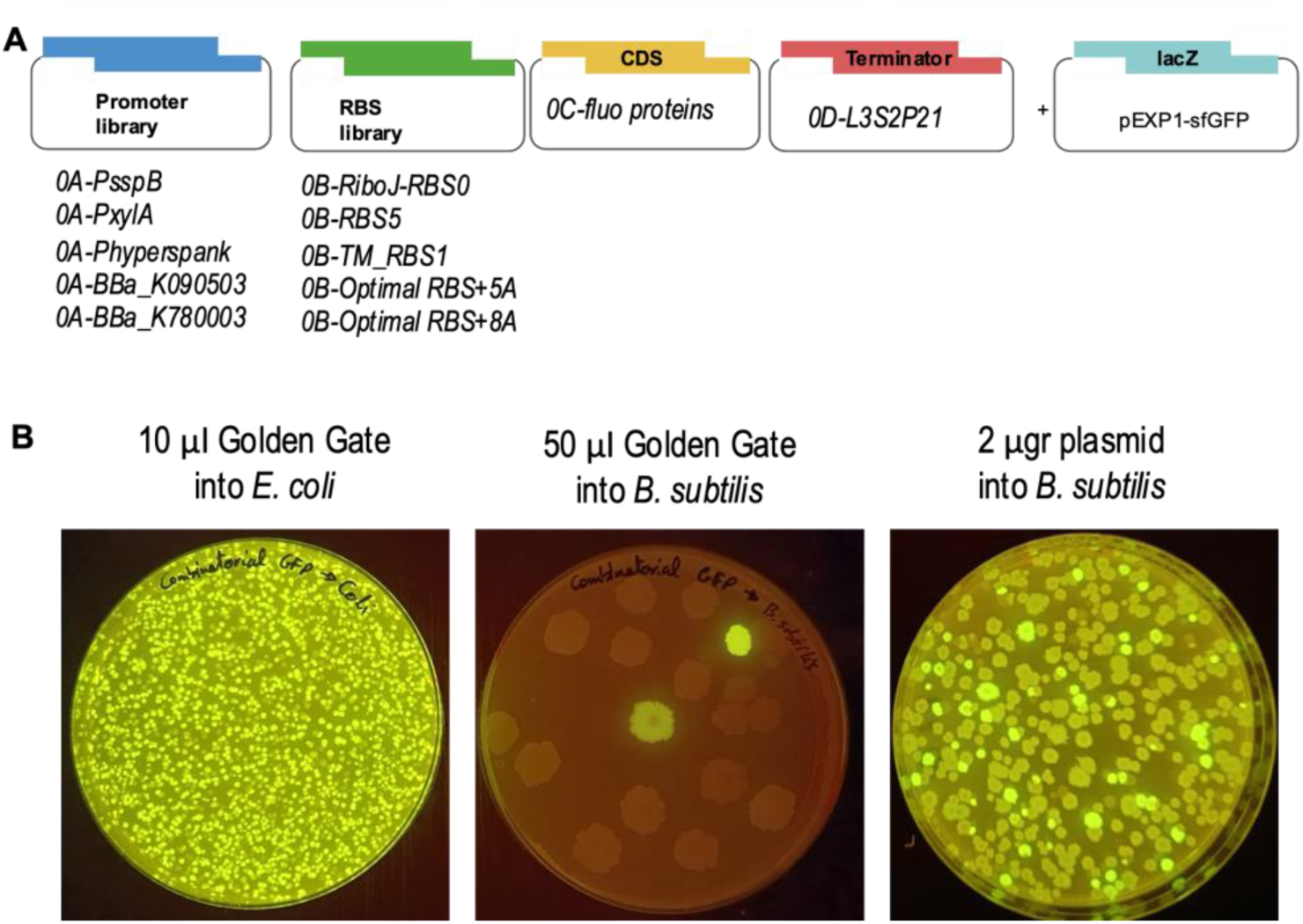
**Combinatorial assemblies into *B. subtilis***. (A) Scheme of the Golden Gate (GG) combinatorial assembly. (B) From a 60 μl volume of a GG reaction, 10 μl were used to transform *E. coli* (left) and 50 μl to transform *B. subtilis* (middle). These efficiencies are contrasted with a transformation of *B. subtilis* using 2 μgr of plasmid DNA purified from a mix culture of 30 different *E. coli* colonies obtained after transformation with GG reaction of a combinatorial assembly.

**Supplementary Figure 7.**
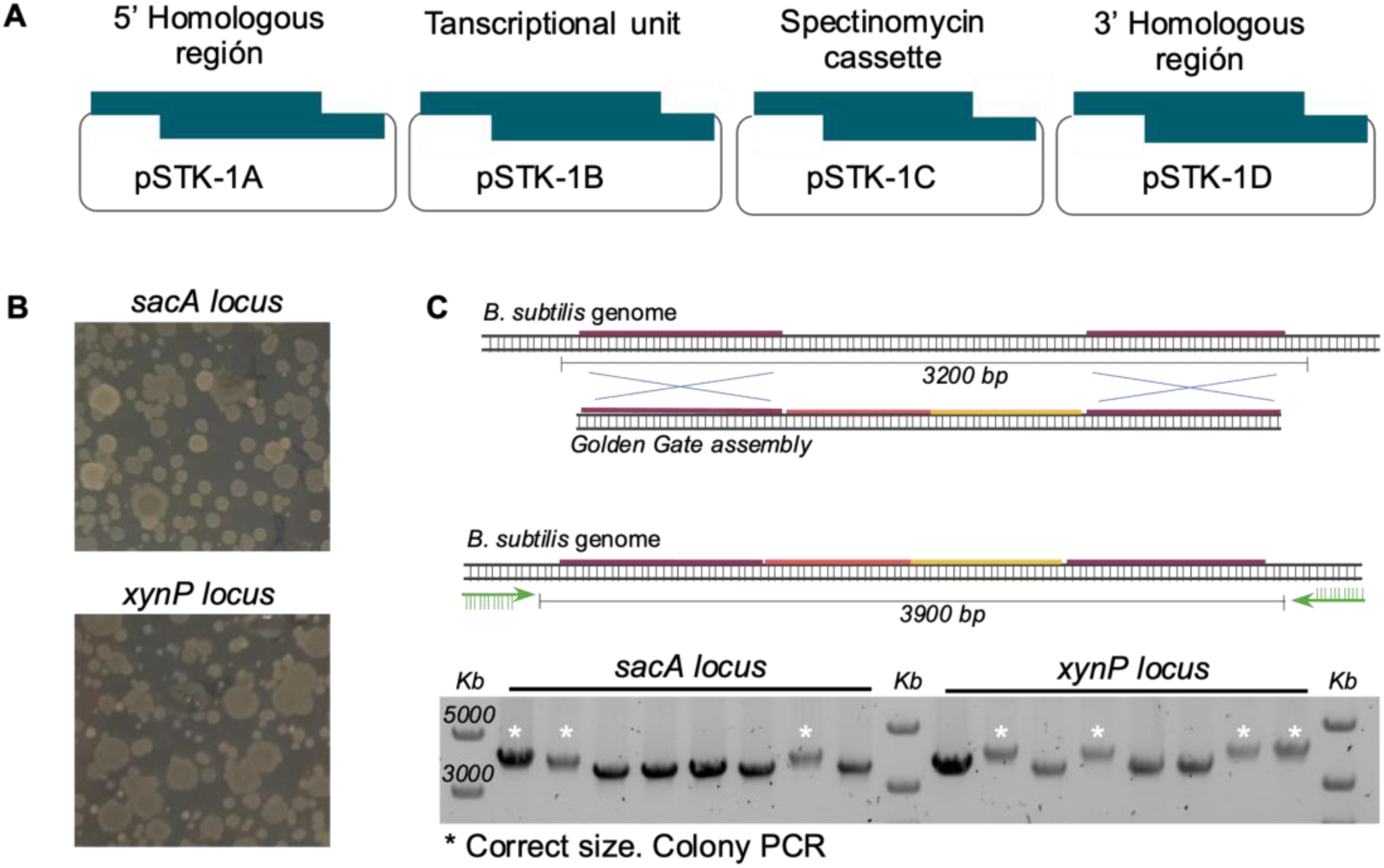
**Genomic integration**. (A) Scheme of the Golden Gate (GG) assembly without destination vector. (B*) B. subtilis* colonies obtained after transformation of 50 μl of GG reaction to assemble the four parts required for double recombination and integration without a destination backbone to produce a linear DNA. (C) Schematic of the double recombination and primers position outside the recombination site to confirm the integration by colony PCR of transformed *B. subtilis* colonies and plated in LB agar supplemented with 100 μg/ml of spectinomycin.

**Supplementary Figure 8.**
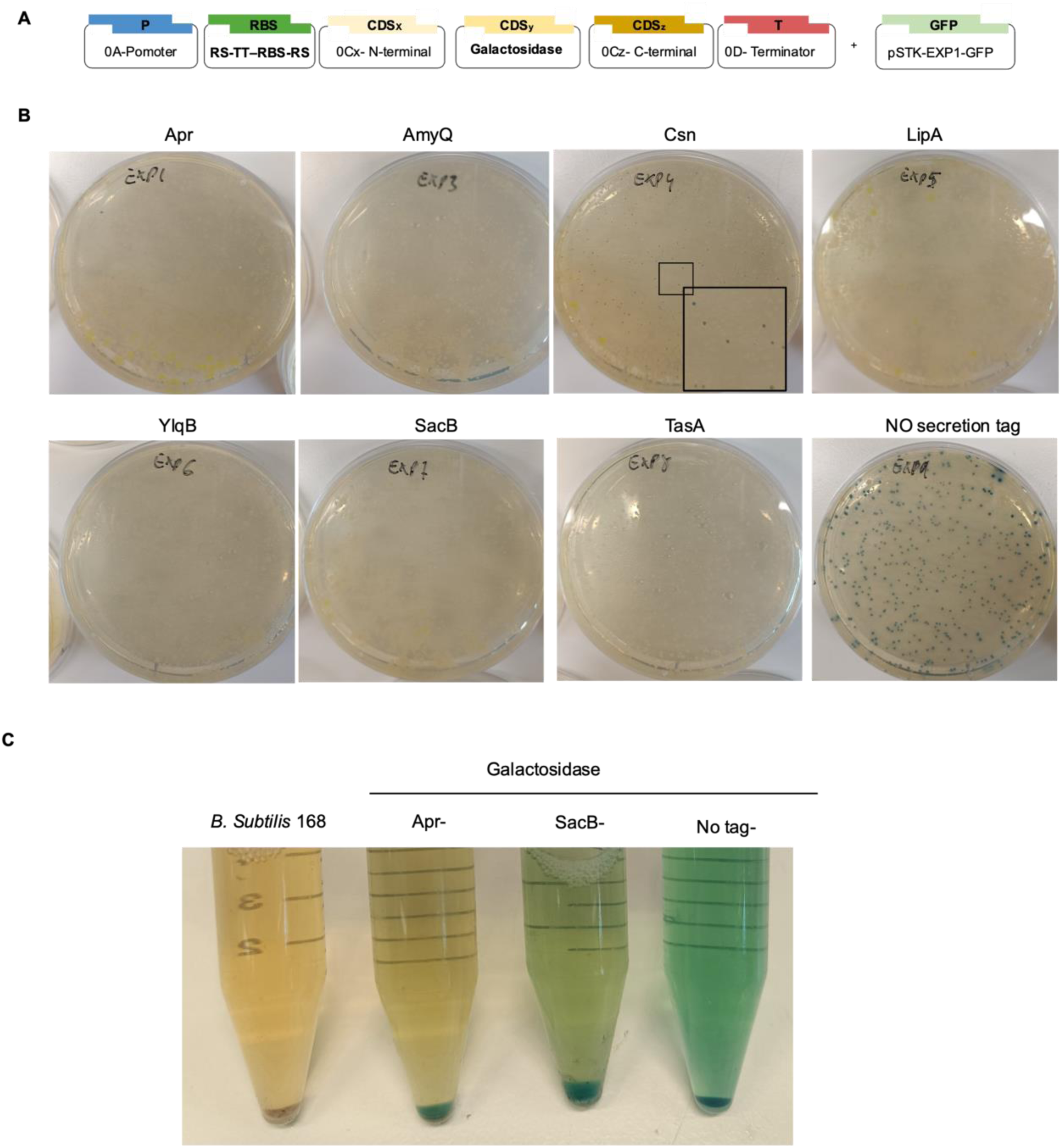
***E. coli* expression blocking device (EEBD)**. (A) Scheme of the Golden Gate (GG) assembly. (B) Images of *E. coli* transformation after 48 h and 37°C incubation. Bacteria were transformed with a GG master mix aliquoted before adding the secretion tag parts to reduce the influence of pipetting errors and were thermocycled together. All the secretion tags were toxic for *E. coli*, obtaining only white colonies (incorrect assemblies) and only Csn-tagged Galactosidase reaction produced tiny blue colonies. (C) Galactosidase activity of Apr and SacB secretion tags compared with the absence of tag shows that the non-classical secretion system is more efficient.

**Supplementary Figure 10.**
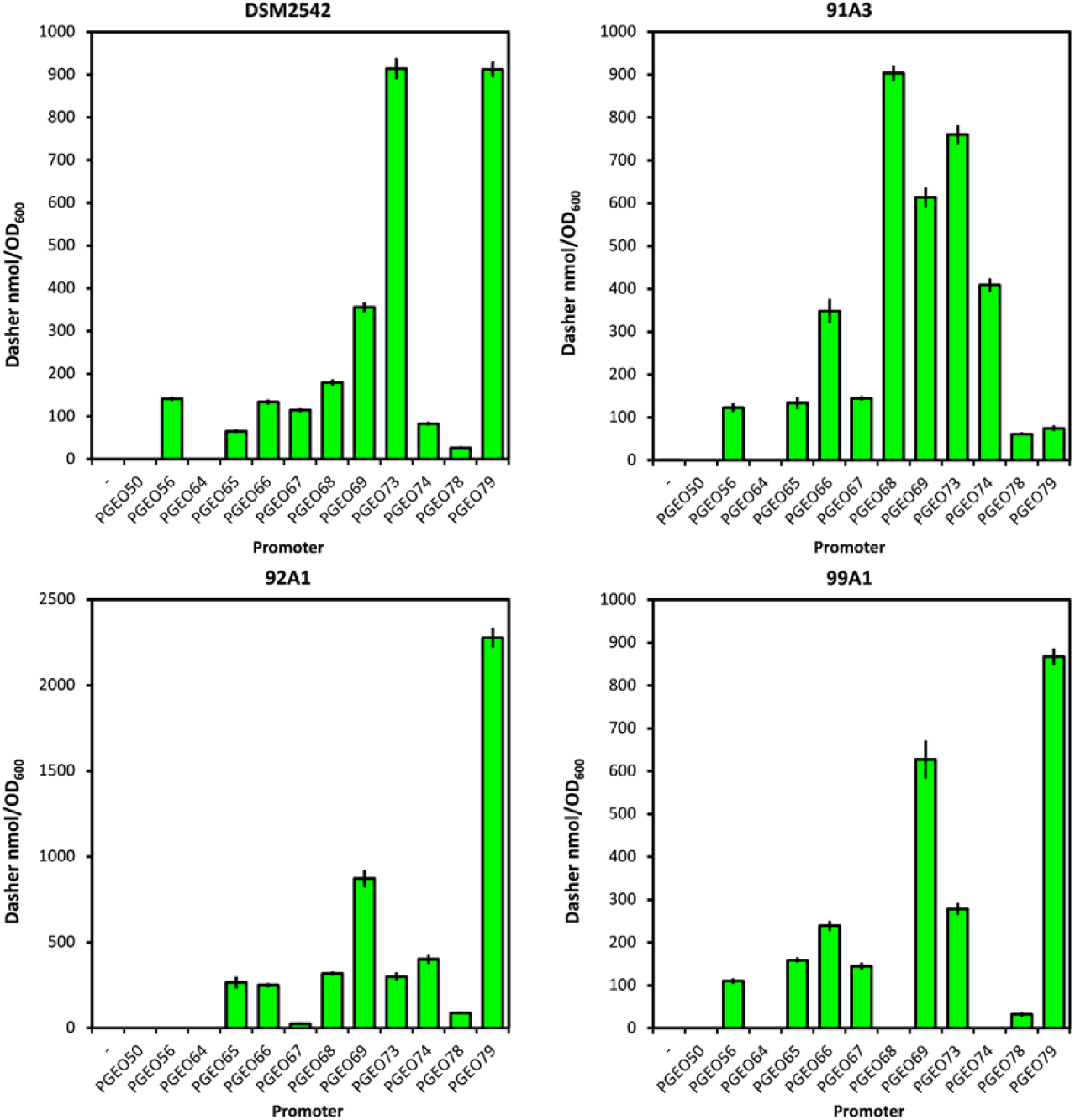
Characterisation of putative *Geobacillus* promoters in different *Geobacillus* species. (DSM2542- *Parageobacillus thermoglucosidasius*, 91A3- *G. subterraneus* strain-K, 99A1- *P. teobii* DSM14590T, 92A1- *G. uzenensis* DSM13551). Promoters were assembled into a pGeoEXP vector with a native RBS from promoter PGEO79, Dasher-GFP as a reporter and a S718 terminator. Transformed cells with their respective reporter constructs were grown in triplicate in 5 ml of mLB for 16 h at 55°C, 200 rpm. Fluorescent intensity (Excitation = 477 nm, emission = 522 nm, Gain 44. Z-position 19131) readings were made and normalised to OD600. Error bars represent standard deviation of three biological repeats.

**Supplementary figure 11.**
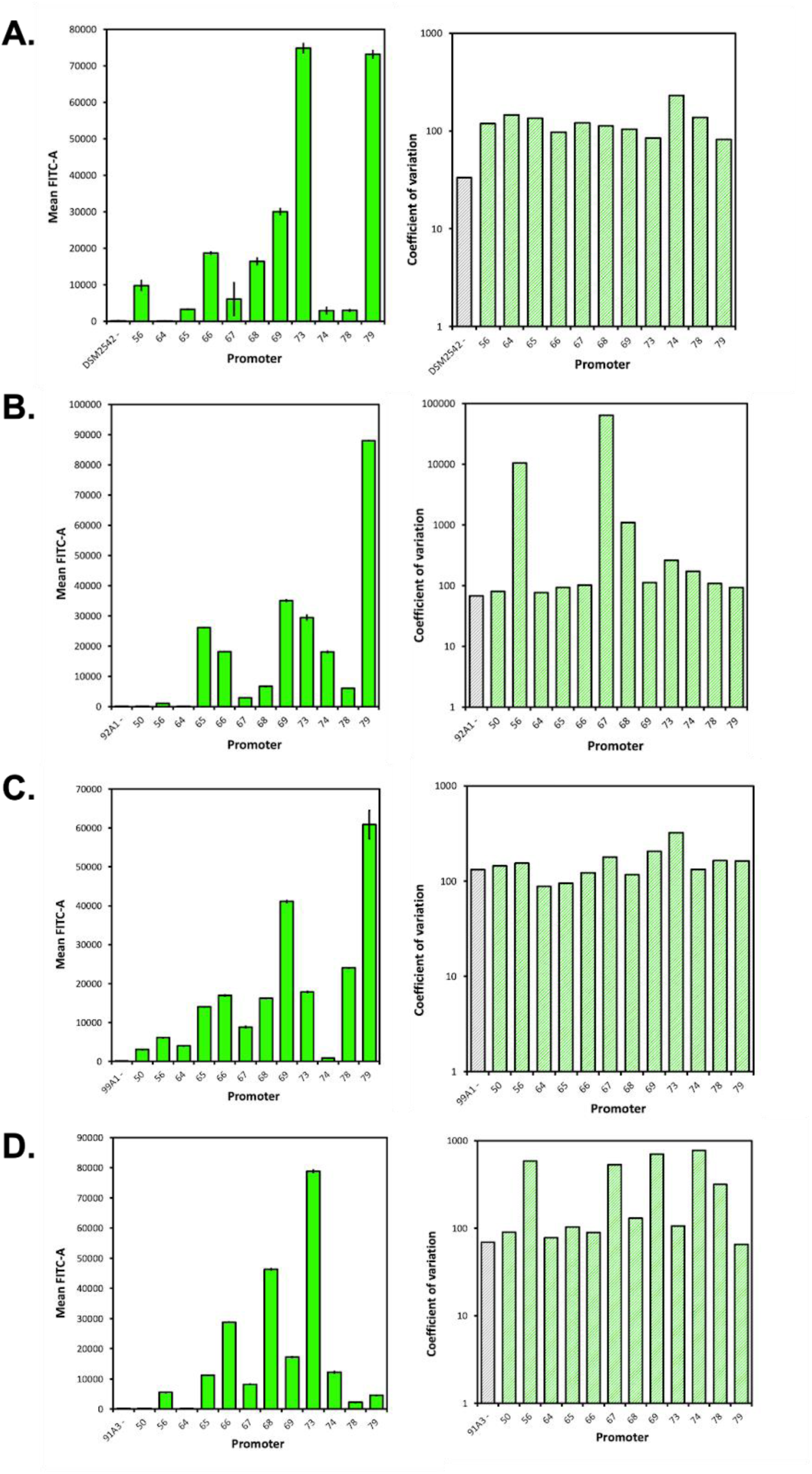
Flow cytometry characterisation of *Geobacillus* promoters in different *Geobacillus* species with a strong *Geobacillus* RBS (PGEO79RBS), with the genetic insulator RiboJ (DSM2542- *Parageobacillus thermoglucosidasius*, 91A3- *G. subterraneus* strain-K, 99A1- *P. teobii* DSM14590T, 92A1- *G. uzenensis* DSM13551). Dasher-GFP was used as a reporter. Cells transformed with their respective reporter construct were grown overnight for 16h in 5 ml of mLB at 55°C at 200 rpm. Cultures were then analysed using a BD FACS Canto II. Dasher-GFP constructs were excited with a 488 nm laser and fluorescence detected using a FITC-A (300 V) fluorophore (Forward scatter = 100, side scatter = 300, threshold = 1,000 FSC). 100,000 events were read for each sample, with three biological repeats, giving a total of 300,000 events per construct tested. Bars indicate the mean fluorescence for each of the samples tested. Error bars indicate standard deviation of three biological repeats. Coefficient of variation (CV) was calculated by dividing population mean by population standard deviation. Data analysis was performed using FlowJo 10.9.0.

**Supplementary Table 1.**
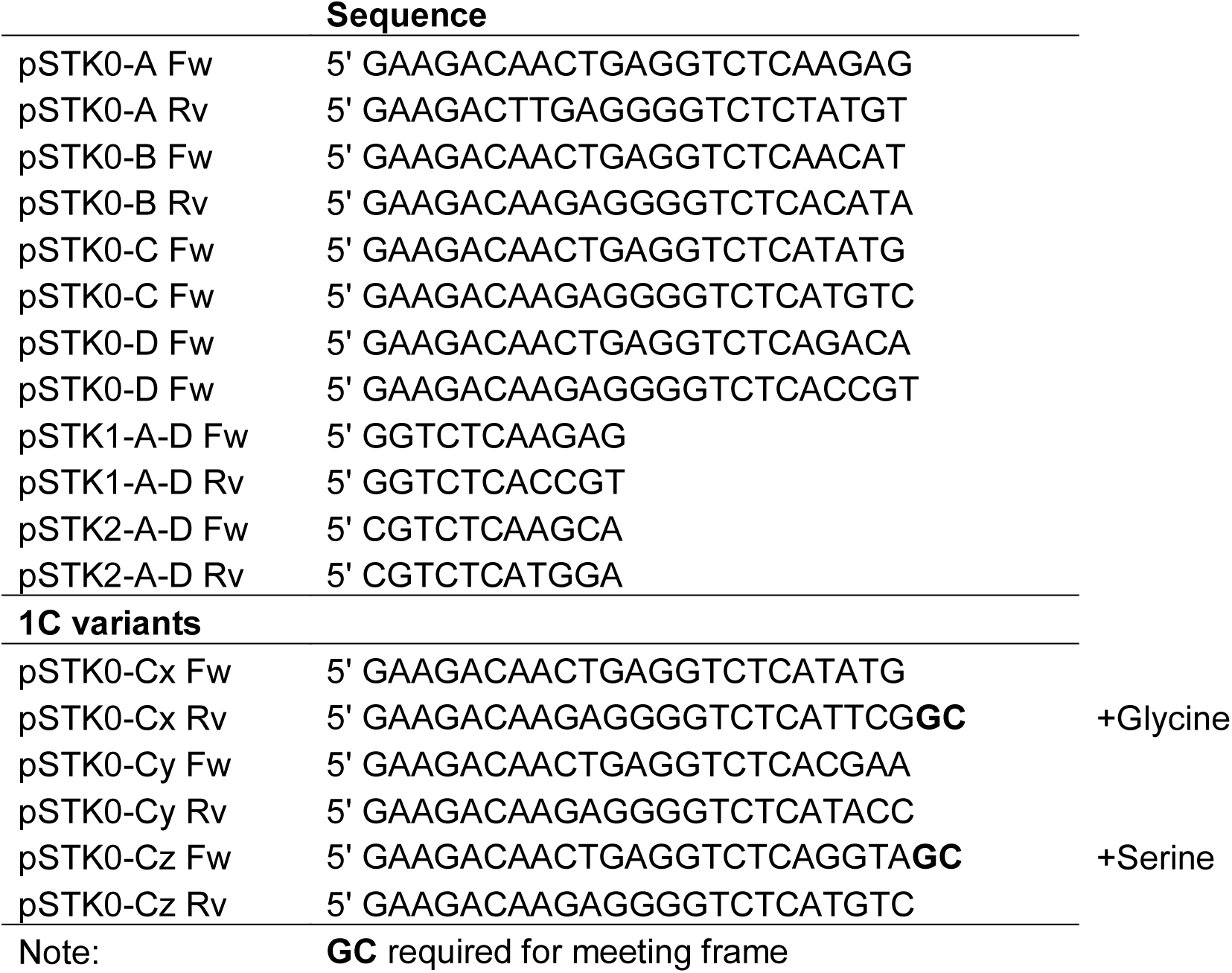
Primer prefixes required to clone parts into STK syntax.

**Supplementary Table 2.**
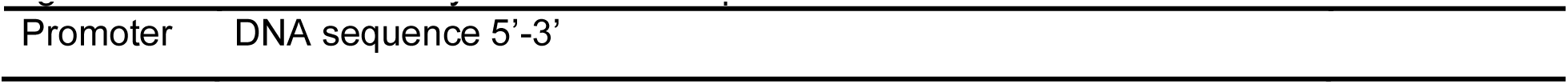
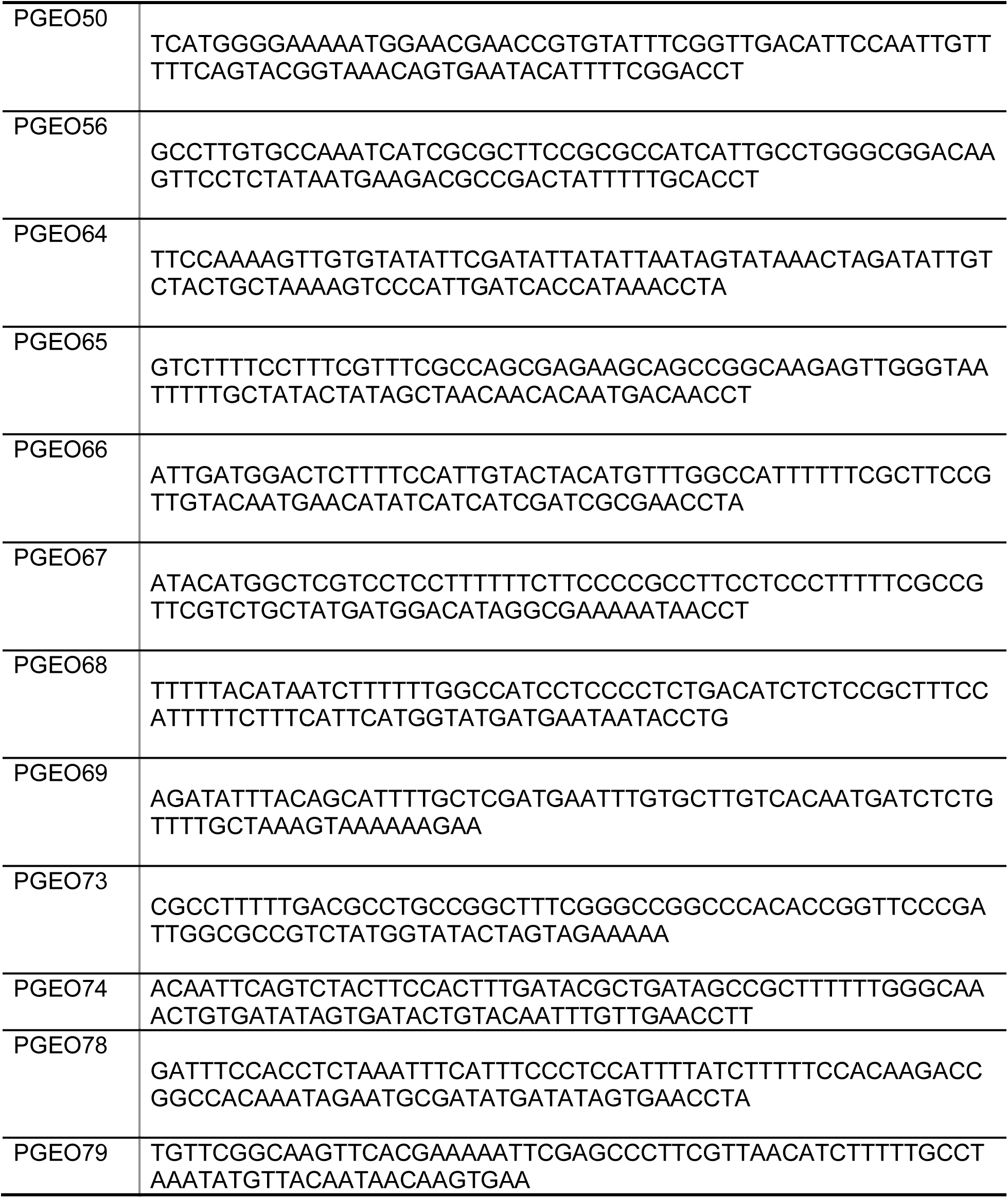
Minimised *Geobacillus* promoters used for STK-GeoToolBox. Promoters from Gilman *et al*. (2019) were modified by removing DNA encoding putative 5’UTR region to allow addition of synthetic RBS sequence.

**Supplementary Table 3.**
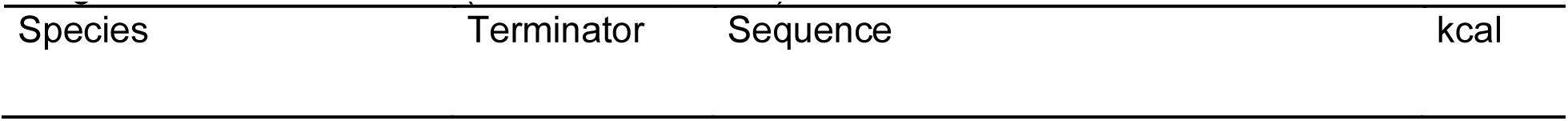
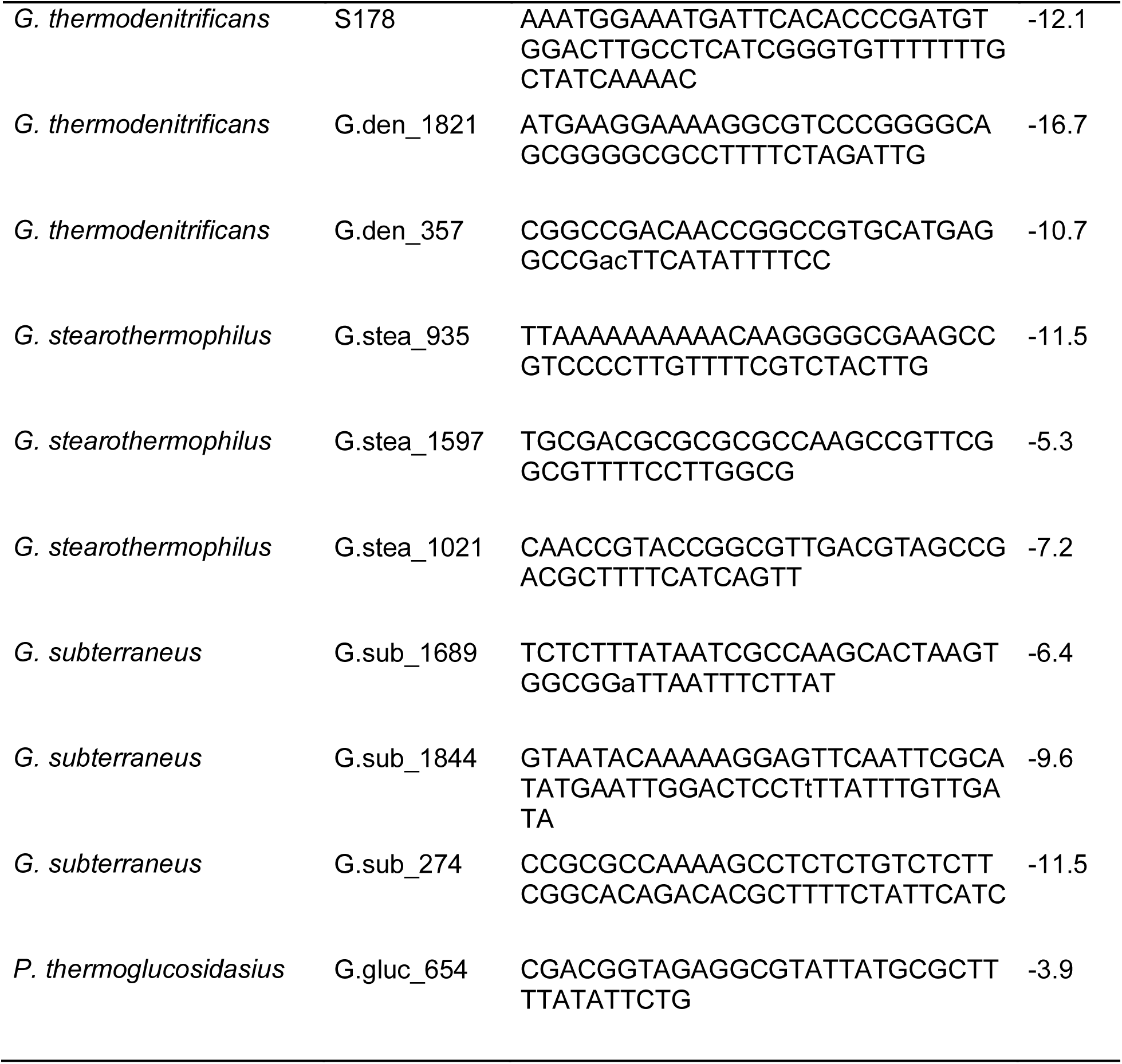
Rho-independent terminators identified in different *Geobacillus* species using the ARNold database (Naville et al., 2011).

**Supplementary Table 4.**
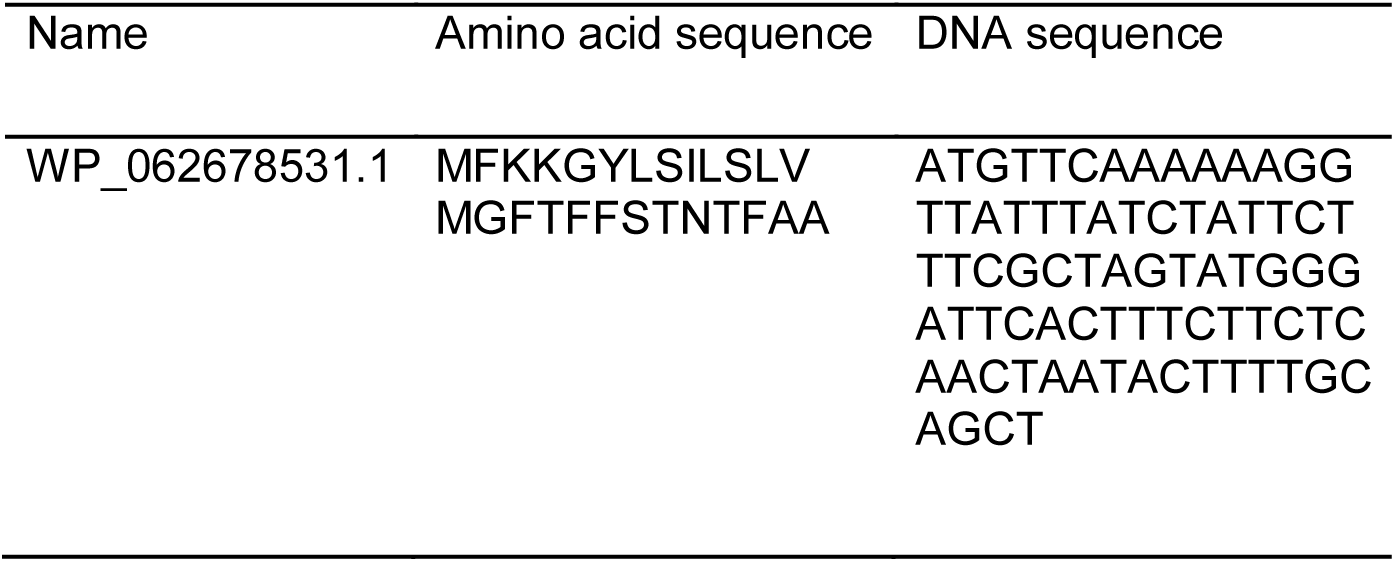
Sec/SPI Signal peptide SPWP_062678531.1 from *P. thermoglucosidasius* DSM2542.

**Supplementary table 5.**
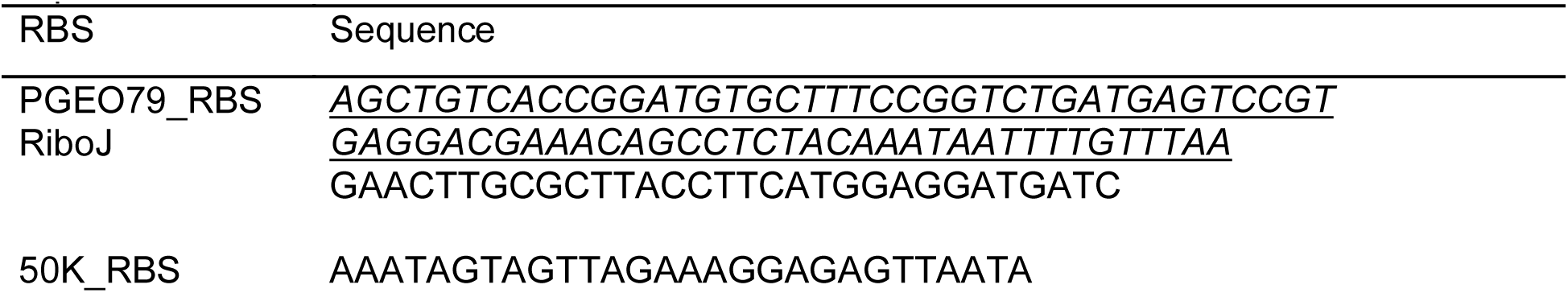
RBS sequences used in this study. Putative RBS from PGEO79 was used for promoter characterisation in different *Geobacillus* and *Parageobacillus* species. Synthetic RBS sequence for Dasher-GFP was designed using the RBS calculator at a translation initiation rate set to 50,000 units(Salis, 2011). Underlined sequence represents the RiboJ sequence.

**Supplementary Table 5.**
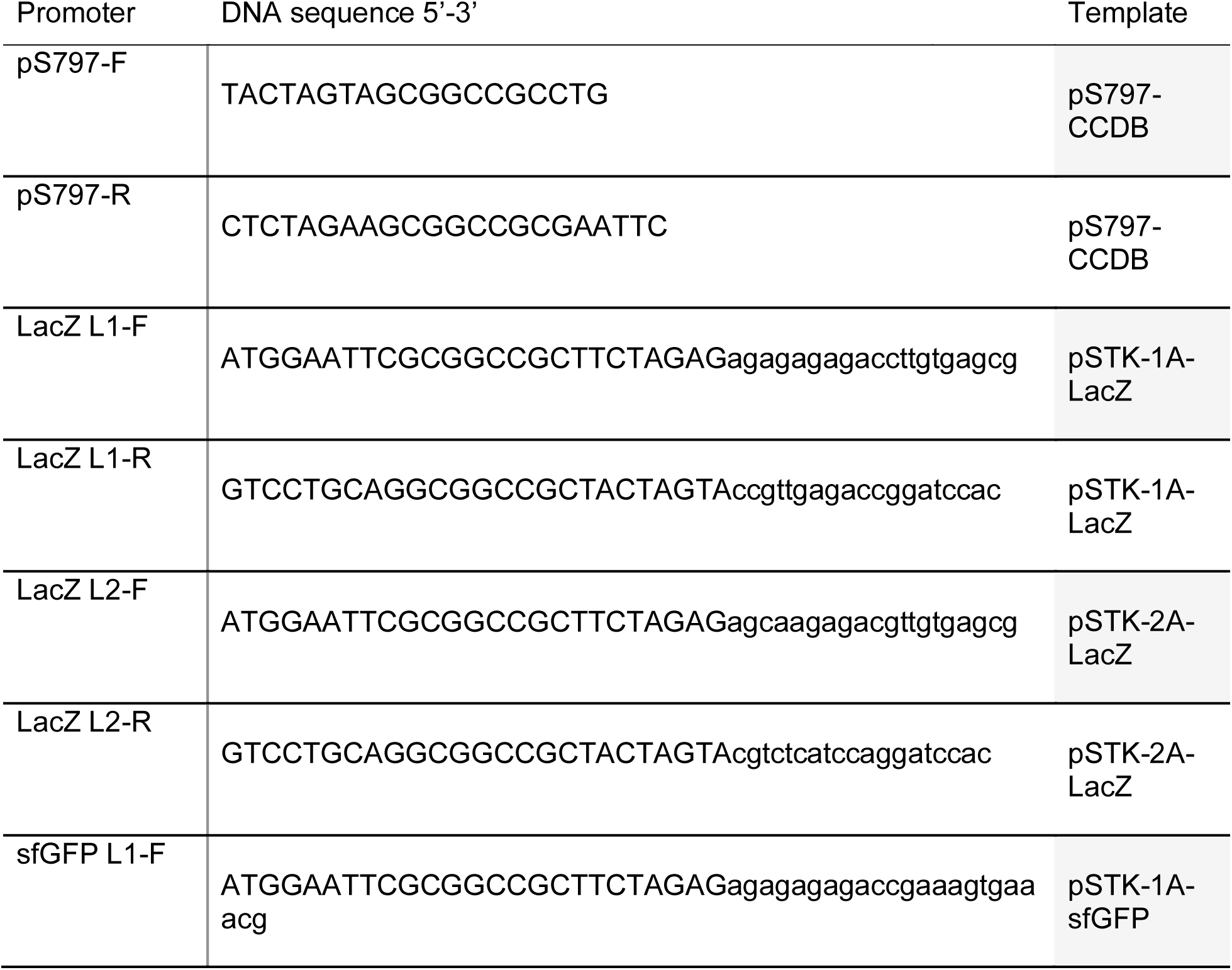
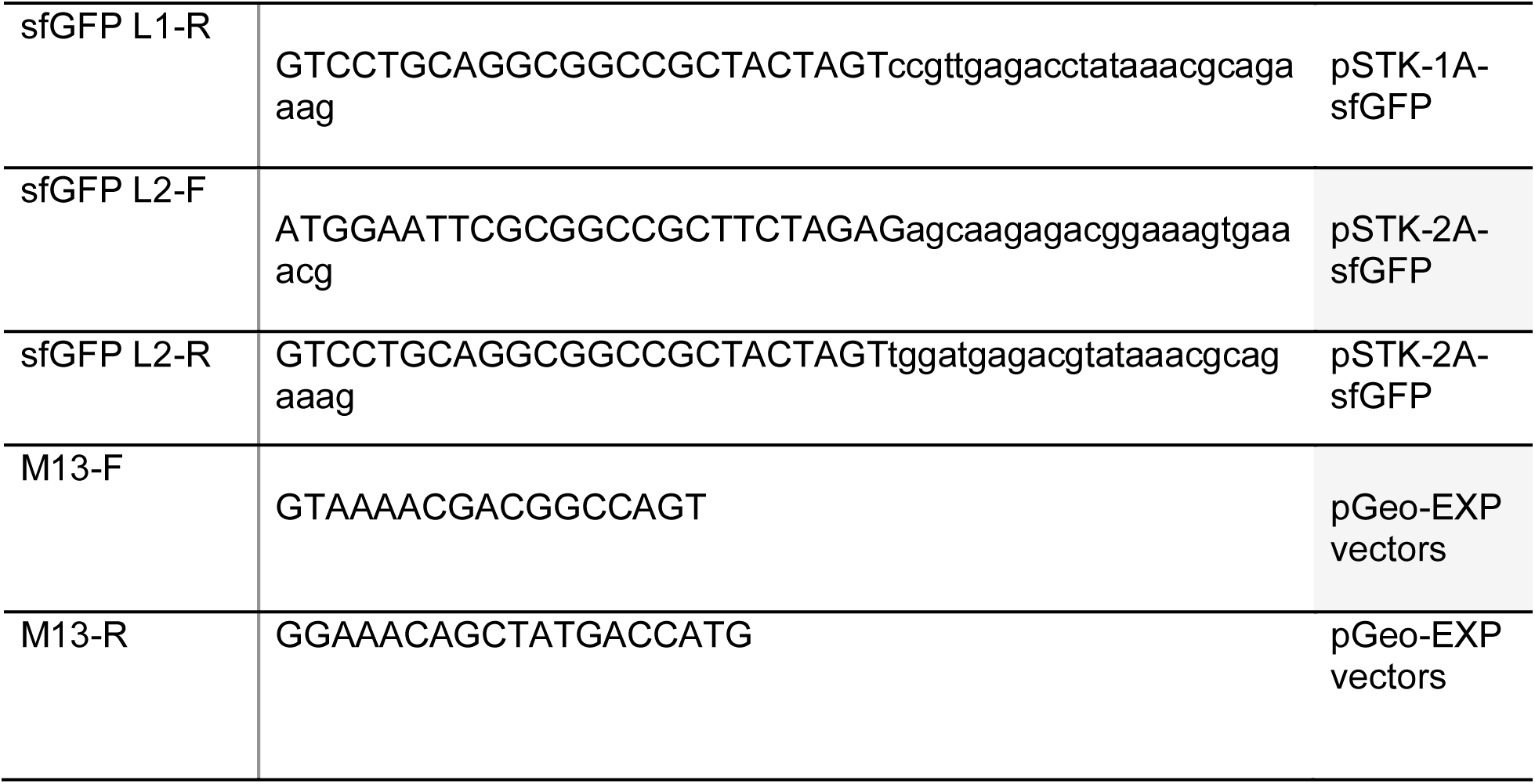
Primers used in this study. F = forward. R = reverse.

## Notes

### Competing Interest Statement

The authors have declared no competing interest.

